# Tissue homeostasis in sponges: quantitative analysis of cell proliferation and apoptosis

**DOI:** 10.1101/2021.10.31.466669

**Authors:** Nikolai P. Melnikov, Fyodor V. Bolshakov, Veronika S. Frolova, Ksenia V. Skorentseva, Alexander V. Ereskovsky, Alina A. Saidova, Andrey I. Lavrov

**Affiliations:** Department of Invertebrate Zoology, Biological Faculty, Lomonosov Moscow State University, Moscow, Russia; Pertsov White Sea Biological Station, Biological Faculty, Lomonosov Moscow State University, Moscow, Russia; Department of Embryology, Biological Faculty, Lomonosov Moscow State University, Moscow, Russia; Department of Cell Biology and Histology, Biological Faculty, Lomonosov Moscow State University, Moscow, Russia; Institut Méditerranéen de Biodiversité et d’Ecologie Marine et Continentale (IMBE), Aix Marseille University, CNRS, IRD, Station Marine d’Endoume, Avignon University, Marseille, France; Department of Embryology, Faculty of Biology, Saint-Petersburg State University, Saint-Petersburg, Russia; Koltzov Institute of Developmental Biology of Russian Academy of Sciences, Moscow, Russia; Department of Cell Biotechnology, Center of Experimental Embryology and Reproductive Biotechnology, Moscow, Russia

**Keywords:** cell turnover, cell proliferation, apoptosis, Porifera, Calcarea, Demospongiae

## Abstract

**Background:** Tissues of multicellular animals are maintained due to a tight balance between cell proliferation and programmed cell death. Phylum Porifera is an early branching group of metazoans essential to understanding the key mechanisms of tissue homeostasis. This paper is dedicated to the comparative analysis of proliferation and apoptosis in intact tissues of two sponges belonging to distinct Porifera lineages, *Halisarca dujardinii* (class Demospongiae) and *Leucosolenia variabilis* (class Calcarea).

**Results:** Labeled nucleotides EdU and anti-phosphorylated histone 3 antibodies reveal a considerable number of cycling cells in intact tissues of both species. The main type of cycling cells are choanocytes - flagellated cells of the aquiferous system. The rate of proliferation remains constant in areas containing choanoderm. Cell cycle distribution assessed by the quantitative DNA stain reveals the classic cell cycle distribution curve. During EdU pulse-chase experiments conducted in *H. dujardinii*, the contribution of the choanocytes to the total amount of EdU-positive cells decreases, while contribution of the mesohyl cells increases. These findings could indicate that the proliferation of the choanocytes is not solely limited to the renewal of the choanoderm, and that choanocytes may participate in the general cell turnover through migration. The number of apoptotic cells in intact tissues of both species is insignificant. *In vivo* studies in both species with TMRE and CellEvent Caspase-3/7 indicate that apoptosis might be independent of mitochondrial outer membrane permeabilization.

**Conclusions:** A combination of confocal laser scanning microscopy and flow cytometry provides a quantitative description of cell turnover in intact sponge tissues. Intact tissues of *H. dujardinii* (Demospongiae) and *L. variabilis* (Calcarea) are highly proliferative, indicating either high rates of growth or cell turnover. Although the number of apoptotic cells is low, apoptosis could still be involved in the regular cell turnover.

## Introduction

Morphogenetic processes underlying the development, growth and regeneration of multicellular animals have long been issues of scientific interest. These phenomena seem to be closely related to a process of physiological regeneration, also known as cell turnover^1,2^. Cell turnover consists of three key elements: 1) the elimination of old, non-functional cells by programmed cell death; 2) the proliferation of somatic stem cells; and 3) the differentiation of newly generated cells along with their integration with preexisting tissue^3^. Physiological regeneration through tissue renewal is an essential process for multicellular animals, as it allows reparation of minor defects and tissue damage occurring during normal functioning^1,4^. Abnormalities of tissue homeostasis are associated with numerous malfunctions, including oncological and degenerative disorders^4,5^.

Sponges (phylum Porifera) are an early-branching metazoan group comprising aquatic, mostly filter-feeding, sedentary organisms^6^. These animals differ substantially from other metazoans in their body organization, lacking tissues and organ systems characteristic of other animals^7,8^. The central anatomical structure in a typical sponge body is its aquiferous system, a complex network of canals and chambers through which the sponge pumps surrounding water for food and oxygen uptake^9^. The rest of the sponge body is composed of the mesohyl, a tissue consisting of an extensive extracellular matrix and numerous wandering cells of different type and function^9^.

Sponge tissues appear to be less specialized than tissues of Eumetazoa. In particular, sponge tissues are multifunctional and highly plastic, as their cells show broad capabilities of transdifferentiation into other cell types^10,11^. The tissue plasticity underlays many aspects of sponge biology, participating in constant reorganization of the aquiferous system^10^, sexual and asexual reproduction^8^, regeneration^12–17^ and even movement^18–20^.

At the same time, sponges lack a single category of somatic stem cells. Choanocytes (flagellated cells of the aquiferous system) and some amoeboid cells of the mesohyl (e.g., archaeocytes of Demospongiae) are both considered to be stem cells in sponges^8,21,22^. It has been shown for several species that choanocytes and archaeocytes proliferate in intact tissues, (trans)differentiate into other cell types and take part in regeneration and gametogenesis^8,14,21,23–25^. These same cells also express some stem cell specific genes, such as members of the germline multipotency program^23,26–28^. Yet, choanocytes are usually interpreted as a minor component of the sponge stem cell system, i.e., the stock of archaeocyte progeny that is able to transform back to archaeocytes^9,23^.

For proper cell turnover, cell proliferation should be balanced by programmed cell death. The most common path of programmed cell death in intact animal tissue is apoptosis^3,29–32^. However, immunohistochemical studies reveal few apoptotic cells in sponges under the steady-state condition^33^. Most of these cells are amoeboid cells located in the mesohyl, with presumably apoptotic choanocytes being extremely rare^33^. On the other hand, some sponges seem to get rid of certain cell types (choanocytes, archaeocytes, lophocytes, spherulous and granular cells) discarding them into the environment through the aquiferous system^33–35^. These expelled cells show various signs of degradation, but the nature of this cell shedding remains obscure^33^.

Considering these peculiarities surrounding tissue organization, sponges can be promising objects for studies regarding the origin and evolution of physiological regeneration mechanisms in metazoans. Therefore, the aim of this study is to provide comparative and quantitative analysis of cell proliferation and apoptosis in intact tissues of two sponge species, *Halisarca dujardinii* Johnston, 1842 (Demospongiae) and *Leucosolenia variabilis* Haeckel, 1870 (Calcarea). These species belong to different Porifera classes and represent principally different types of aquiferous system organization^12,36^. Using combined approaches of confocal laser scanning microscopy and flow cytometry, we have identified proliferating and apoptotic cells in sponge tissue while evaluating their number and distribution throughout the sponge body. Altogether, this data complements the concept of tissue dynamics in these basal metazoans.

## Results

### Cell proliferation in intact sponge tissues: quantitative analysis and spatial distribution

*Halisarca dujardinii* (class Demospongiae) is a thick globular sponge with 1-2 oscular tubes and a leuconoid aquiferous system (Figure 1A, B). Its body is subdivided into two regions, the ectosome and the endosome (Figure 1C). The ectosome represents a superficial region of the body devoid of choanocyte chambers. The external parts of the T-shaped exopinacocytes are underlined with a thick layer of diffuse collagen that contains rare spherulous cells (Figure 1C). A dense layer of amoeboid cells (e.g., spherulous and granular cells) and cell bodies of the exopinacocytes are located beneath the diffuse collagen layer (Figure 1C). The internal region of a sponge, the endosome, contains large, tubular choanocyte chambers surrounded by the mesohyl (Figure 1C, D). The mesohyl contains numerous cells (e.g., archaeocytes, lophocytes, spherulous, vacuolar and granular cells) and occupies a significant volume of the endosome (Figure 1C). The structure of the aquiferous system is similar in regions of the endosome adjacent to the oscular tube, and at some distance (approximately half the body radius) from it (Figure 2A, B). The oscular tube is a principally different region of a sponge, representing a thin-walled outgrowth composed mainly of exo- and endopinacoderm (Figure 2C). A thin layer of mesohyl lies between the exo- and endopinacoderm. Choanocyte chambers are absent.

**Figure 1.**
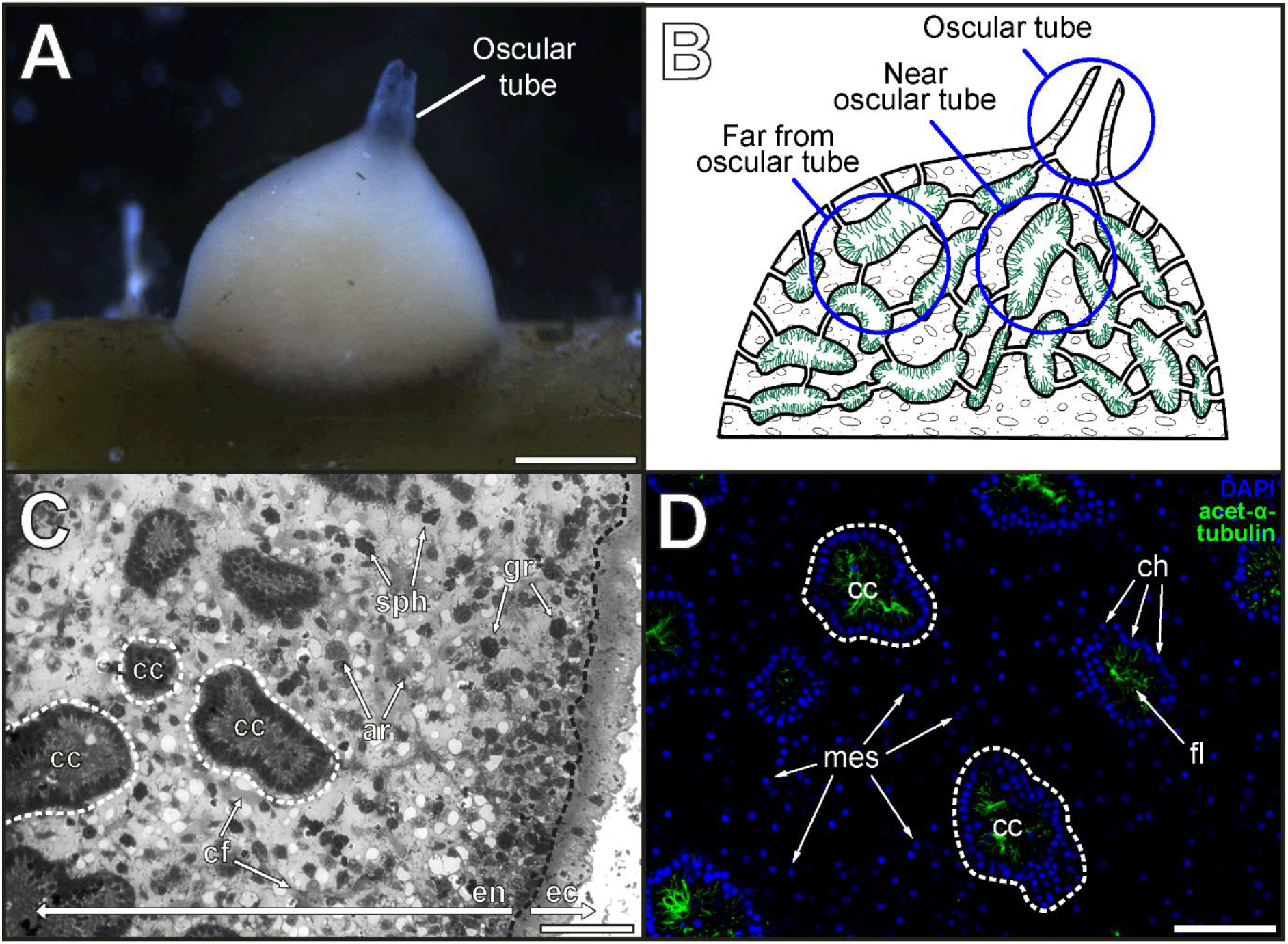
Anatomy and histology of *Halisarca dujardinii*. A – general appearance; B – schematic drawing of the aquiferous system representing regions of interest; C – section through the endosome and ectosome using light microscopy; D – area of the endosome distant from the oscular tube, CLSM (maximum intensity projection of several focal planes); blue color – DAPI, cell nuclei; green – acetylated α-tubulin, flagella. Black dotted line marks the border between the ecto- and endosome. White dotted lines mark the borders of several choanocyte chambers. ar – archaeocyte, cc – choanocyte chamber, cf – collagen fiber, en – endosome, ec – ectosome, fl – flagella, gr – granular cell, mes – nucleus of mesohyl cell, sph – spherulous cell. Scale bars: A – 1 mm, C, D – 50 μm.

**Figure 2.**
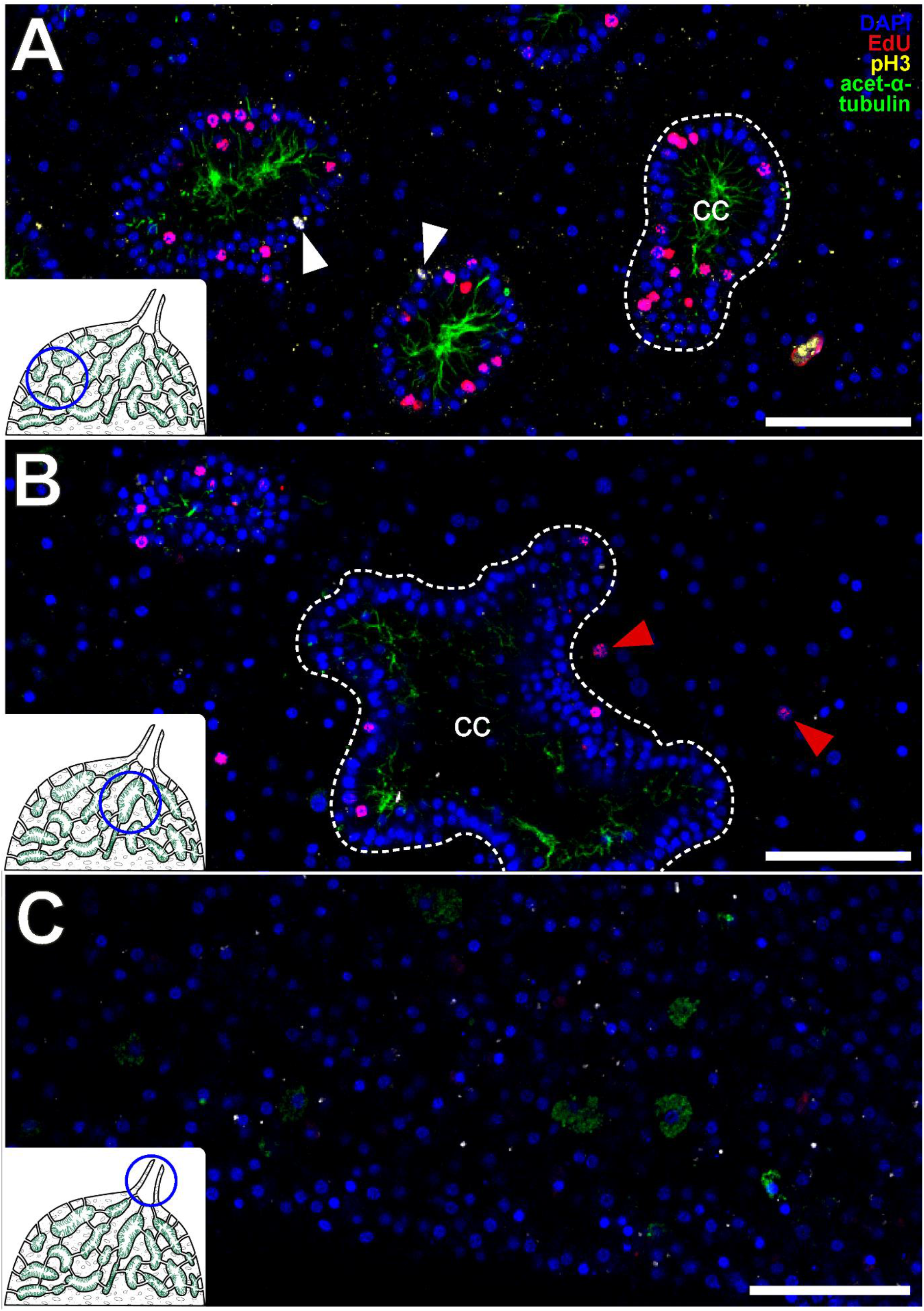
Cell proliferation in different body regions of *Halisarca dujardinii*, CLSM. A - endosome distant from the oscular tube; B - endosome adjacent to the oscular tube; C – the oscular tube. All images are maximum intensity projections of several focal planes. Insets schematically show position of the studied regions in the sponge body. Blue color – DAPI, cell nuclei; red – EdU, nuclei of DNA synthesizing cells; yellow – pH3, chromatin of late G2/M cells; green – acetylated α-tubulin, flagella. Dotted lines mark the borders of choanocyte chambers. White arrowheads mark cells in the M-phase of the cell cycle; red arrowheads mark EdU-positive cells outside the choanocyte chambers. cc - choanocyte chambers (outlined by the dotted lines). Scale bars: A, B, C - 50 μm.

To evaluate the number and distribution of proliferating cells in intact tissues of *Halisarca dujardinii*, we have utilized two markers: 5-ethynyl-2’-deoxyuridine (EdU) and Ser10-phosphorylated histone 3 (pH3) monoclonal antibodies. After 12 hours of treatment with 200 μM EdU, intact tissues of *H. dujardinii* contained numerous EdU-positive nuclei belonging to a population of cycling cells (Figure 2A, B). Few cells showed nuclei labeled with pH3-antibodies, representing a fraction of cycling cells in the late G2 or M phases. Even fewer were pH3-positive cells containing EdU in their nuclei. The endosome adjacent to the oscular tube and the endosome distant from the oscular tube displayed similar fraction of proliferating cells, containing 8.44±3.08% and 8.10±1.64% of EdU-positive cells, respectively. Each of the beforementioned regions also had 0.11±0.07% of pH3-positive cells (Figure 3A, B; Table 1). Dunn test shows no significant differences between these regions (p-values>0.9999 for both markers). Both regions also showed a similar distribution of proliferating cells; most of the EdU- and pH3-positive nuclei (approximately 90% of all labeled nuclei) were located in the choanocyte chambers and belonged to choanocytes, while EdU- and pH3-positive nuclei in the mesohyl are few (Figures 2A, B; 3C, D; Table 1).

**Figure 3.**
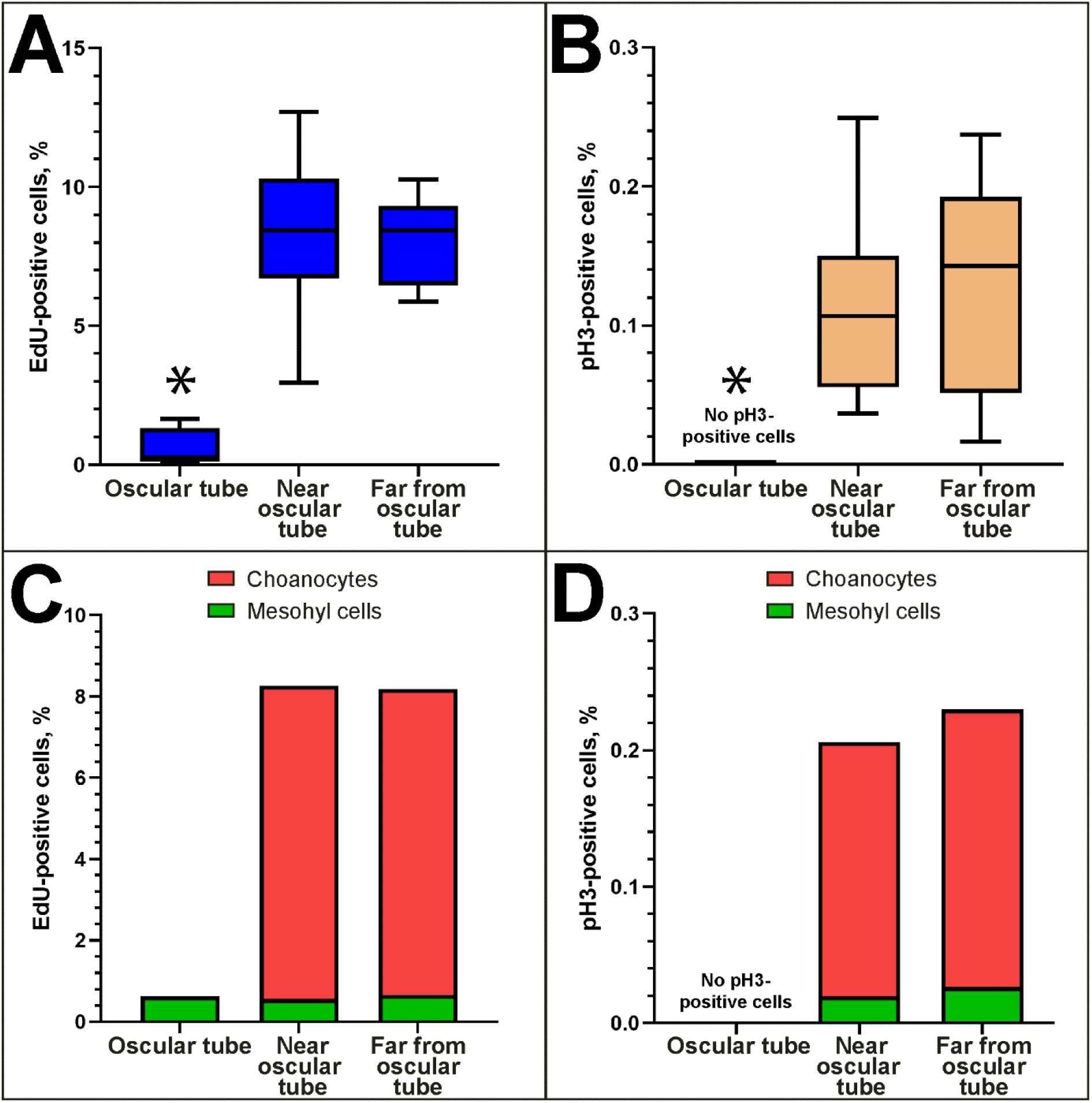
Quantitative analysis of cell proliferation in intact tissues of *Halisarca dujardinii*. A, B - fractions of EdU- (A) and pH3-positive (B) cells in different body regions. Data is shown with median values (thick horizontal lines), interquartile ranges (boxes), total ranges (whiskers) and outliers (dots). Asterisks mark significantly different regions of the body. C, D – fractions (mean values) of choanocytes/mesohyl cells among the total number of EdU- (C) and pH3-positive (D) cells.

**Table 1.**
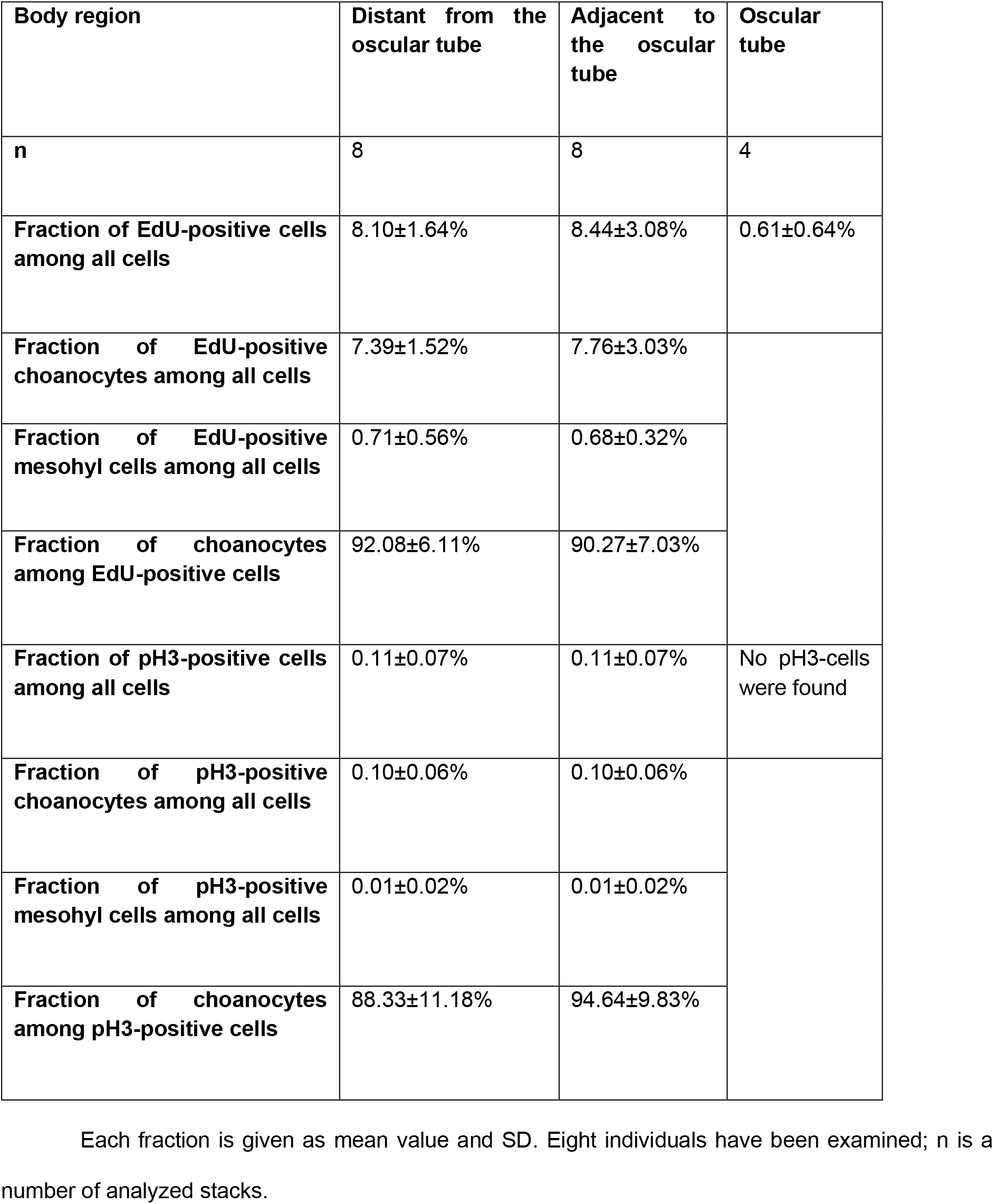
Fractions of proliferating cells in intact tissues of *Halisarca dujardinii*, CLSM.

There were 0.61±0.64% of EdU-positive cells and no pH3-positive cells in the oscular tube (Figures 2C; 3A, B; Table 1). All EdU-positive cells belonged to the mesohyl/pinacoderm, as this region is devoid of choanocyte chambers (Figure 3C). The fraction of EdU-positive cells in the oscular tube is significantly lower than the total fraction of EdU-positive cells in the regions of the endosome (p<0.05 in each comparison) (Figure 3A). The fraction of EdU-positive cells in the oscular tube showed no difference with regards to the fraction of EdU-positive mesohyl cells in the regions of the endosome (p-values>0.9999 in each comparison) (Figure 3C).

*Leucosolenia variabilis* (class Calcarea) is an asconoid sponge; its body is a plexus of branching and anastomosing thin-walled hollow tubes forming the cormus (Figure 4A). The cormus gives rise to the oscular tubes and the blind outgrowths or diverticula. The body wall is composed of the exopinacoderm and choanoderm layers separated by a thin mesohyl layer containing few cells, mostly amoebocytes and sclerocytes (Figure 4C-G). The cell bodies with nuclei of T-shaped exopinacocytes and porocytes are located near the mesohyl cells (Figure 4E). The choanocytes are easily distinguished from other cells as they form a continuous layer of tightly packed pyriform nuclei associated with flagella (Figure 4E-G). The structure of the body wall as described is characteristic of most body regions in L. variabilis: diverticula, cormus tubes and the oscular tubes. In the oscular rim, the most distal part of the oscular tube, the structure of the body wall is altered since choanocytes are substituted by flat endopinacocytes (Figure 4C).

**Figure 4.**
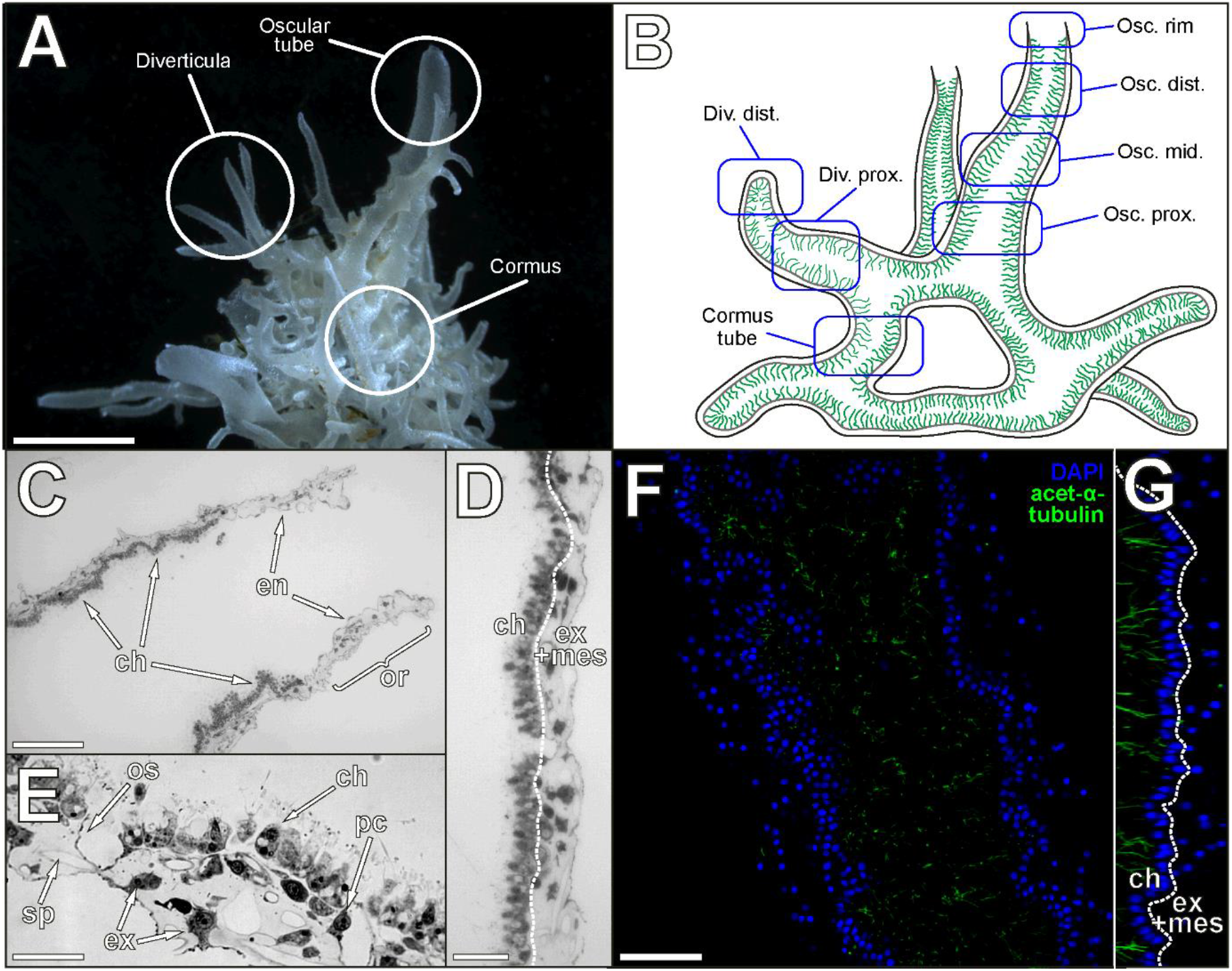
Anatomy and histology of *Leucosolenia variabilis*. A – general appearance; B – schematic drawing of the aquiferous system representing regions of interest; C – longitudinal section through the distal part of the oscular tube (including the oscular rim), light microscopy; D, E – cross sections through the middle part of the oscular tube, light microscopy; F – the proximal part of the divericulum, CLSM (maximum intensity projection of several focal planes); G – orthogonal XZ projection of the same Z-stack, CLSM (maximum intensity projection of several focal planes); blue color – DAPI, cell nuclei; green – acetylated α-tubulin, flagella. Osc. rim – the oscular rim; Osc. dist - distal part of the oscular tube; Osc. mid. - middle part of the oscular tube; Osc. prox. - proximal part of the oscular tube; Div. dist. - distal part of the diverticulum; Div. prox. - proximal part of the diverticulum; Cormus tube - cormus tube. ch – choanoderm; en – endopinacocytes, ex – exopinacocytes; ex+mes – exopinacoderm/mesohyl layer; or – oscular rim; os – ostium; pc – porocyte; sp – spicule. Dotted lines mark the borders between the choanoderm and exopinacoderm/mesohyl layers. Scale bars: A – 0.2 mm; C, D – 50 μm; E – 25 μm; F – 50 μm.

Intact tissues of *L. variabilis* displayed a significant number of cycling cells after 6 hours of treatment with 20 μM EdU and pH3-antibodies labeling (Figure 5A-G). Few pH3-positive cells displayed EdU signal in the nuclei (Figure 5H). Regions containing choanocytes exhibited a high proliferation rate. Most of these regions (*i.e*., cormus tubes, proximal parts of diverticula, oscular tubes) showed a similar fraction of proliferating cells (p-values>0.9999 in each pairwise comparison), each containing about 10% of EdU-positive and 0.6% of pH3-positive cells (Figures 5B-G; 6A, B; Table 2). In the distal parts of the diverticula, the proportion of pH3-positive cells remained the same (0.50±0.20%), while the proportion of EdU-positive cells decreased (6.53±2.08%) (Figure 6A, B; Table 2). The decrease of EdU-positive cells is significant when compared to the distal and middle parts of oscular tubes (p-values=0.0068 and 0.0022, respectively). In other pairwise comparisons, differences were are insignificant, but p-values were slightly higher than 0.05. The oscular rim had the lowest fraction of proliferating cells when compared to other body parts, containing only 0.40±0.96% of EdU-positive and 0.17±0.20% of pH3-positive cells (Figure 6A, B; Table 2). The decrease in the number of EdU labeled cells is significant when compared across other body regions, except for the distal part of the diverticula.

**Figure 5.**
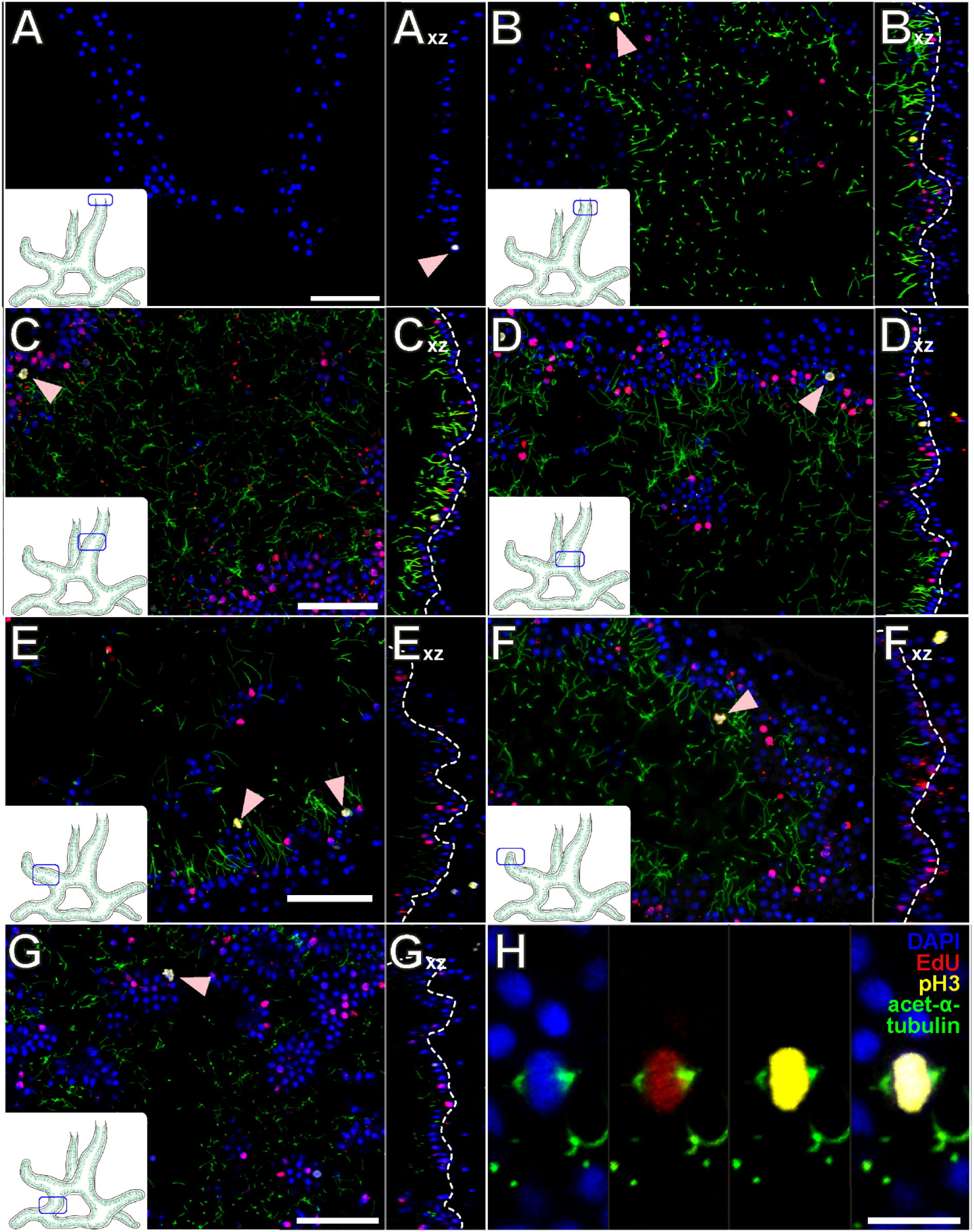
Cell proliferation in different body regions of *Leucosolenia variabilis*, CLSM. A – oscular rim; B – the distal part of an oscular tube; C - the middle part of an oscular tube; D - the proximal part of an oscular tube; E - the proximal part of a diverticulum; F - the distal part of a diverticulum; G - a cormus tube; E – metaphase plate in the choanoderm of the middle part of the oscular tube. Images with the “xz” index represent XZ orthogonal projections of the respective Z-stack. All images are maximum intensity projections of several focal planes. Insets schematically show position of the studied regions in the sponge body. Blue color – DAPI, cell nuclei; red – EdU, nuclei of DNA synthesizing cells; yellow – pH3, chromatin of late G2/M cells; green – acetylated α-tubulin, flagella. Arrowheads mark cells in the M-phase of the cell cycle. Scale bars: A, B, C, D, E, F, G – 50 μm.

**Figure 6.**
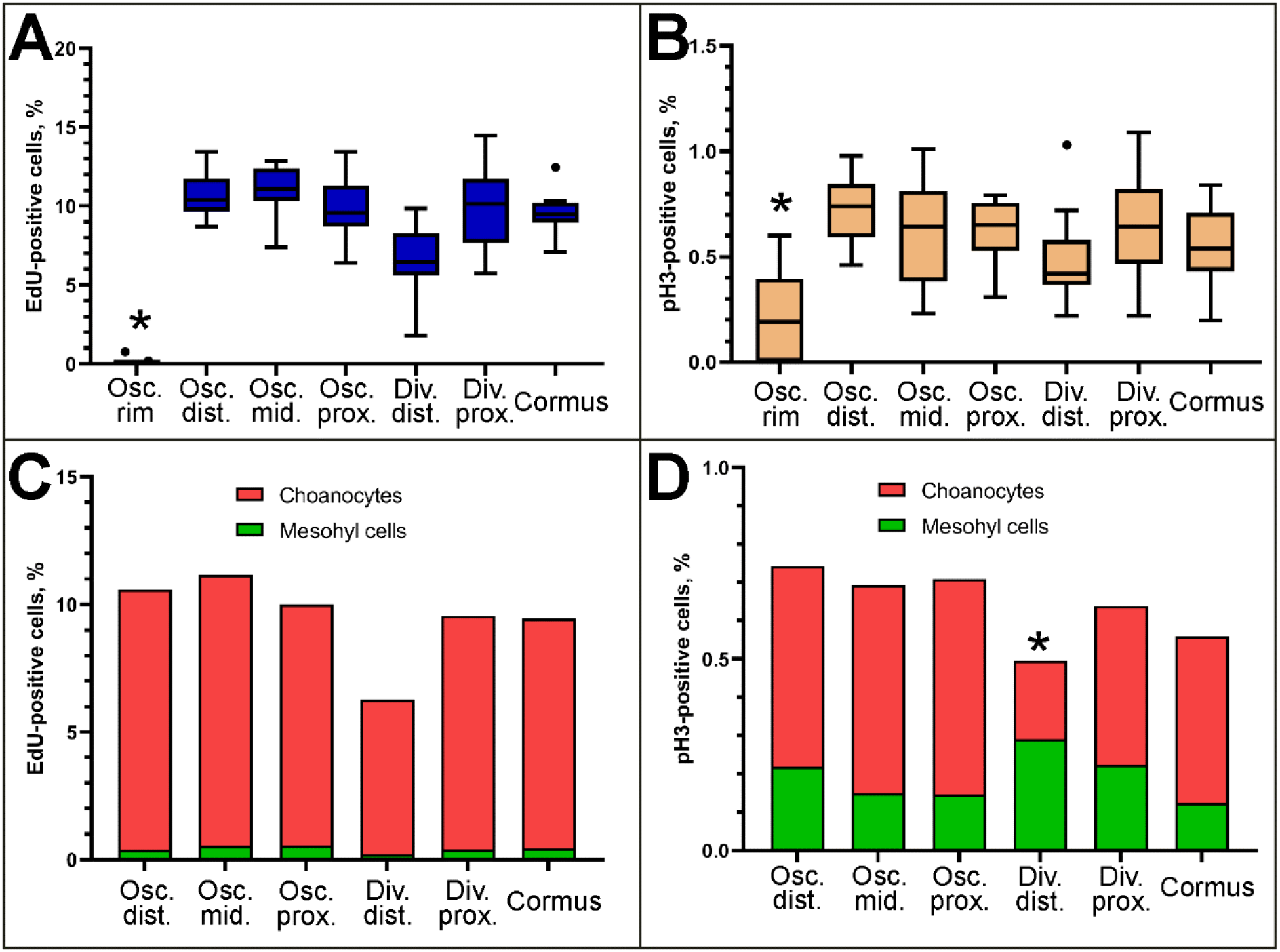
Quantitative analysis of cell proliferation in different body regions of *Leucosolenia variabilis*. A, B - fractions of EdU- (A) and pH3-positive (B) cells in different body regions. Data is shown with median values (thick horizontal lines), interquartile ranges (boxes), total ranges (whiskers) and outliers (dots). C, D – fractions (mean values) of choanocytes/mesohyl cells among the total number of EdU- (C) and pH3-positive (D) cells, respectively. Asterisks mark significantly different regions of the body. Osc. rim - oscular rim; Osc. dist - distal part of an oscular tube; Osc. mid. - middle part of an oscular tube; Osc. prox. - proximal part of an oscular tube; Div. dist. - distal part of a diverticulum; Div. prox. - proximal part of a diverticulum; Cormus - cormus tube.

**Table 2.**
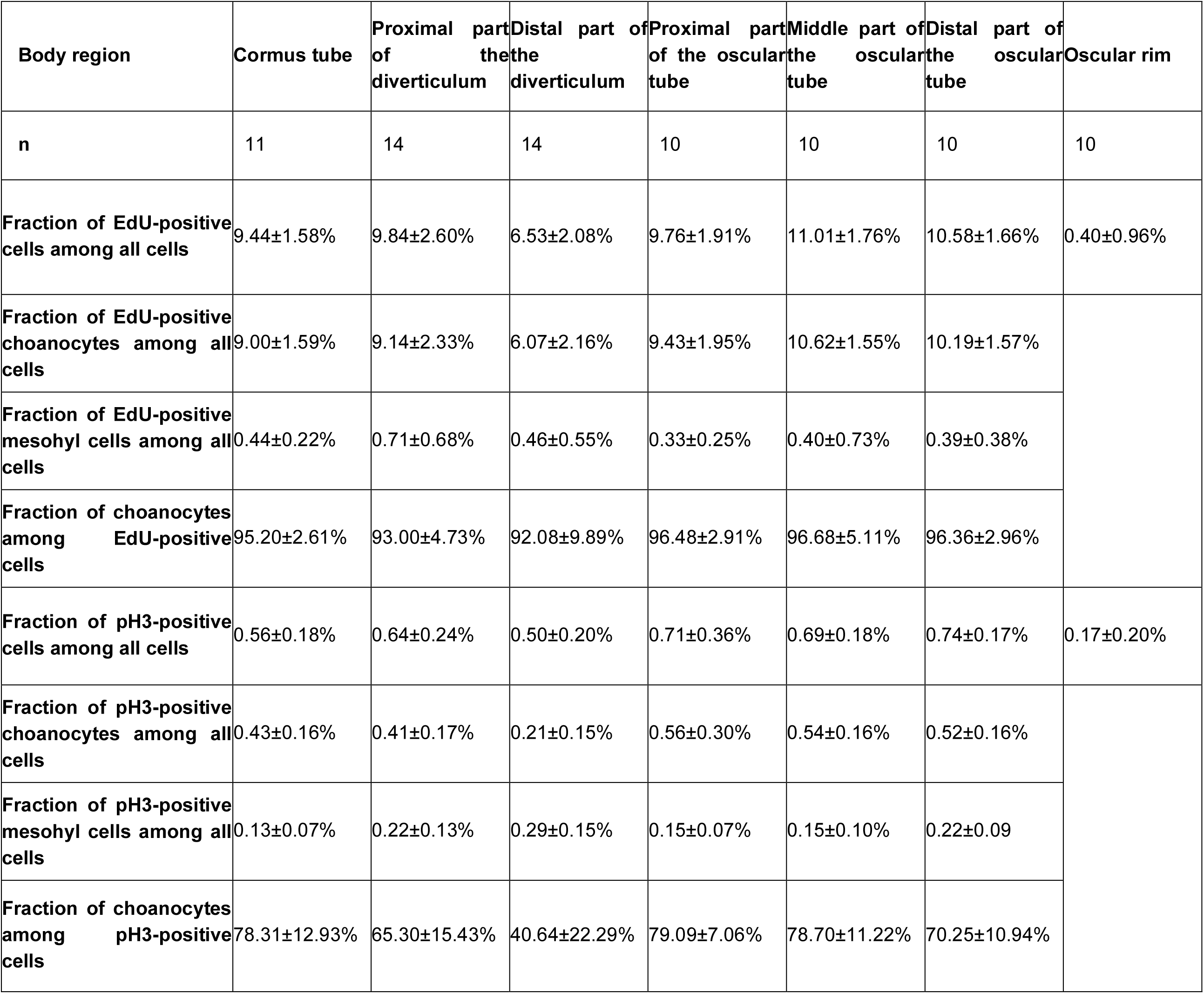
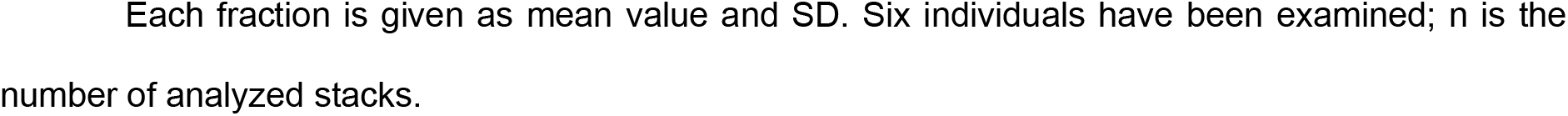
Fractions of proliferating cells in different body regions of *Leucosolenia variabilis*, CLSM.

| Body region | Cormus tube | Proximal part of the diverticulum | Distal part of the diverticulum | Proximal part of the oscular tube | Middle part of the oscular tube | Distal part of the oscular tube | Oscular rim |
| --- | --- | --- | --- | --- | --- | --- | --- |
| n | 11 | 14 | 14 | 10 | 10 | 10 | 10 |
| Fraction of EdU-positive cells among all cells | 9.44±1.58% | 9.84±2.60% | 6.53±2.08% | 9.76±1.91% | 11.01±1.76% | 10.58±1.66% | 0.40±0.96% |
| Fraction of EdU-positive choanocytes among all cells | 9.00±1.59% | 9.14±2.33% | 6.07±2.16% | 9.43±1.95% | 10.62±1.55% | 10.19±1.57% |  |
| Fraction of EdU-positive mesohyl cells among all cells | 0.44±0.22% | 0.71±0.68% | 0.46±0.55% | 0.33±0.25% | 0.40±0.73% | 0.39±0.38% |  |
| Fraction of choanocytes among EdU-positive cells | 95.20±2.61% | 93.00±4.73% | 92.08±9.89% | 96.48±2.91% | 96.68±5.11% | 96.36±2.96% |  |
| Fraction of pH3-positive cells among all cells | 0.56±0.18% | 0.64±0.24% | 0.50±0.20% | 0.71±0.36% | 0.69±0.18% | 0.74±0.17% | 0.17±0.20% |
| Fraction of pH3-positive choanocytes among all cells | 0.43±0.16% | 0.41±0.17% | 0.21±0.15% | 0.56±0.30% | 0.54±0.16% | 0.52±0.16% |  |
| Fraction of pH3-positive mesohyl cells among all cells | 0.13±0.07% | 0.22±0.13% | 0.29±0.15% | 0.15±0.07% | 0.15±0.10% | 0.22±0.09% |  |
| Fraction of choanocytes among pH3-positive cells | 78.31±12.93% | 65.30±15.43% | 40.64±22.29% | 79.09±7.06% | 78.70±11.22% | 70.25±10.94% |  |

1 Each fraction is given as mean value and SD. Six individuals have been examined; n is the 2 number of analyzed stacks.

Most of the EdU- and pH3-positive cells of *L. variabilis* are choanocytes. However, a few of the EdU-positive and pH3-positive cells belonged to the mesohyl/pinacoderm layer (Figures 5A-G; 6C, D). The proportion of choanocytes in the total number of EdU-positive cells remained almost the same (95% on average) in different parts of the oscular tube, cormus and diverticula (Figure 6C; Table 2). With regards to the pH3-positive cells, choanocytes represented approximately 75% of all the late G2/M cells (Figure 6D; Table 2). In the distal parts of the diverticula, their proportion decreased (40.64±22.29%; Figure 6D; Table 2). This decrease is significant in comparison to the cormus, middle and proximal parts of the oscular tube (p=0.0023, 0.0003 and 0.0002, respectively).

### EdU tracking in Halisarca dujardinii under the steady-state condition

Since labeled nucleotides are transferred to daughter cells after mitosis, EdU allows us to track proliferating cells or their progeny. We treated individuals of *H. dujardinii* with 200 μM EdU for 12 hours and moved them into the running seawater aquarium. In the aquarium, the incorporation of EdU ceases and the distribution of EdU-positive cells in intact tissue of *H. dujardinii* gradually changes. Specifically, the contribution by choanocytes to the total amount of EdU-positive cells decreases while contribution by mesohyl cells increases (Figure 7A, B). At the start of the experiment the mesohyl cells accounted for only 7.92±6.11% of the labeled cells; after 7 days their fraction significantly increased, reaching 55.12±23.15% (p=0.0026) (Figure 7C).

**Figure 7.**
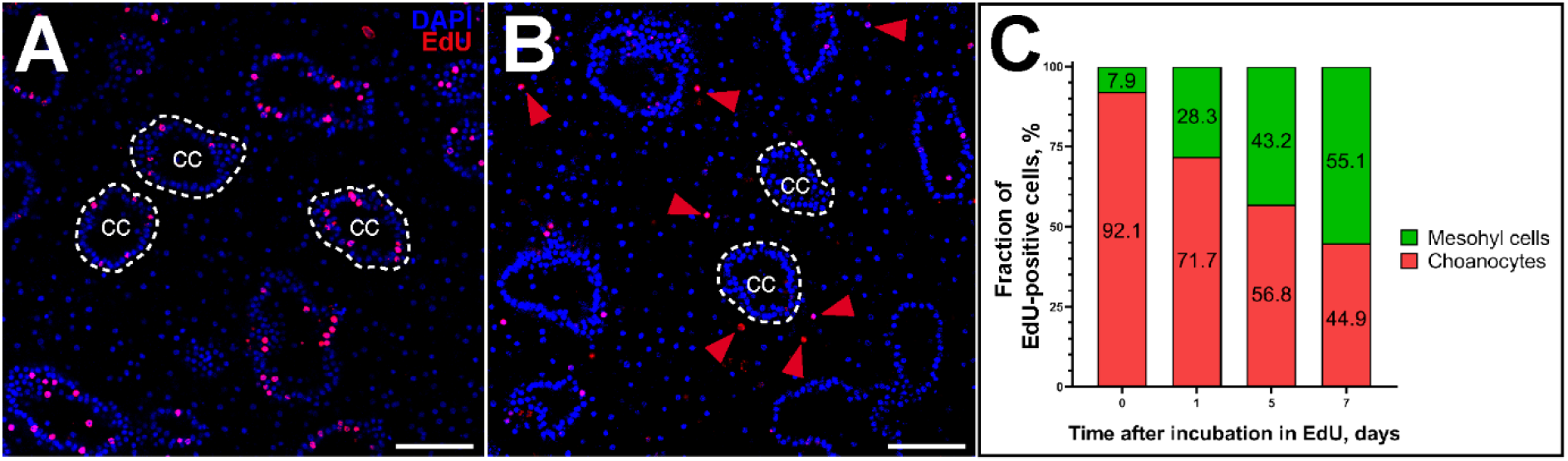
EdU tracking experiment in *Halisarca dujardinii*, CLSM. A - 0 days after the incubation in EdU (maximum intensity projection of several focal planes); B - 7 days after the incubation in EdU (maximum intensity projection of several focal planes); blue color – DAPI, cell nuclei, red – EdU, nuclei of DNA synthesizing cells. C - fractions of labeled choanocytes and mesohyl cells among all EdU-positive cells (mean values). Dotted lines mark the borders of choanocyte chambers. Arrowheads in B mark EdU-positive nuclei in the mesohyl. Scale bars: A, B - 50 μm.

### Cell cycle distribution in cell suspensions of Halisarca dujardinii and Leucosolenia variabilis

Cell cycle distribution was assessed by the quantitative DNA staining of single-cell suspensions obtained via mechanical dissociation in calcium and magnesium-free seawater (CMFSW-E). The major part of *H. dujardinii* cells (38.94±5.90%) formed a diploid G0/G1 population peak (Figure 8A; Table 3). The tetraploid cells in the G2/M phase formed a less distinguishable peak characterized by a 2-fold higher fluorescence intensity than that of the G1/G0 cells (Figure 8A). The G2/M population contained 9.28±1.78% of all cells (Table 3). Cells in the S phase were located between the G0/G1 and G2/M populations and accounted for 20.18±6.14% of the total population (Table 3). The SubG1 population was characterized by lower DNA content than G1/G0 cells (Figure 8A). Scatter plots indicated that the SubG1 population consisted of small and highly granulated objects - possibly, dying cells and debris (Figure 8C).

**Figure 8.**
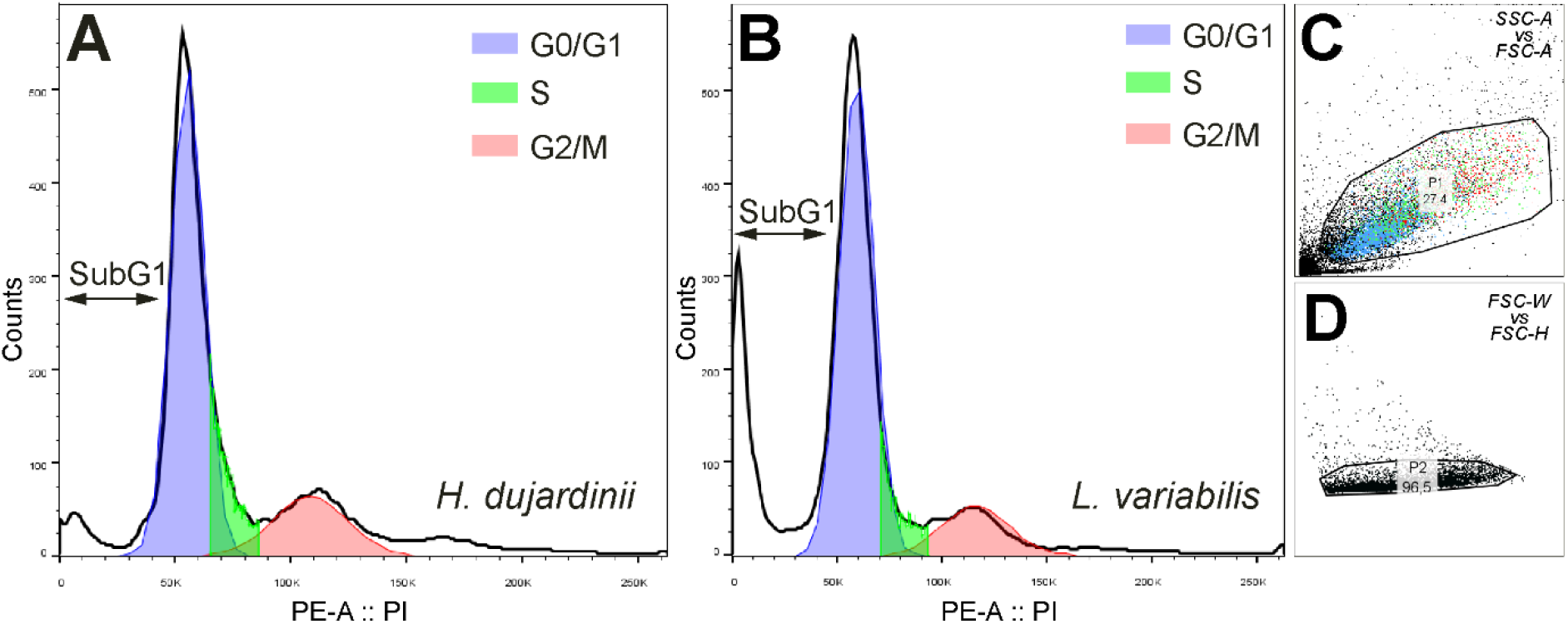
Single-parameter histograms of PI fluorescence (DNA content) in *Halisarca dujardinii* (A) and *Leucosolenia variabilis* (B) cell suspensions; C, D – an example of light-scatter primary gating to distinguish single cells in a suspension of *L. variabilis*. G0/G1 (red) are diploid cells, G2/M (blue) are tetraploid cells, S (green) are DNA-synthesizing cells. SubG1 cells are cells with fragmented DNA.

**Table 3.** Fractions of cells in different phases of the cell cycle, flow cytometry.

| Species | N | SubG1 | G0/G1 | S | G2/M |
| --- | --- | --- | --- | --- | --- |
| *Halisarca dujardinii* | 5 | 23.41±12.22 % | 38.94±5.9% | 20.18±6.14% | 9.28±1.78% |
| *Leucosolenia variabilis* | 5 | 26.42±3.43% | 53.88±2.59% | 12.58±0.76% | 4.63±1.79% |

4 Each proportion is given as mean value and SD.

CMFSW-E suspensions of *L. variabilis* did not provide reliable data: they contained too many SubG1 and too few G2/M cells to obtain an accurate cell cycle distribution histogram and were ultimately excluded from further analysis. We therefore used suspensions made from pre-fixed tissue of *L. variabilis*. This method provided improved resolution: suspensions contained fewer SubG1 cells and showed well-distinguishable G0/G1 and G2/M peaks. The general appearance of the histogram was similar to *H. dujardinii* (Figure 8B). The proportion of G0/G1 cells was 53.88±2.59%, 12.58±0.76% for S cells and 4.63±1.79% for G2/M cells (Table 3).

### Investigation of apoptotic activity

To study the apoptosis, individuals of *H. dujardinii* and *L. variabilis* were vitally stained with fluorescent substrate for effector caspases, CellEvent, for two hours. Subsequent CLSM studies revealed an insignificant number of apoptotic cells in intact tissues of the sponges (Figure 9A, D). Most of these cells exhibited a weak CellEvent fluorescence and were differentiated from autofluorescent cells by nuclear localization of the signal characteristic of CellEvent (Figure 9B-F). In *L. variabilis*, the proportion of CellEvent-positive cells did not exceed 1% in any of the studied body regions (Table 4). Both choanocytes and mesohyl cells were present among the apoptotic cells. The endosome of *H. dujardinii* showed even less apoptotic activity; for every 6-9 thousand normal cells, there were usually 1-2 CellEvent-positive ones (Table 4).

**Figure 9.**
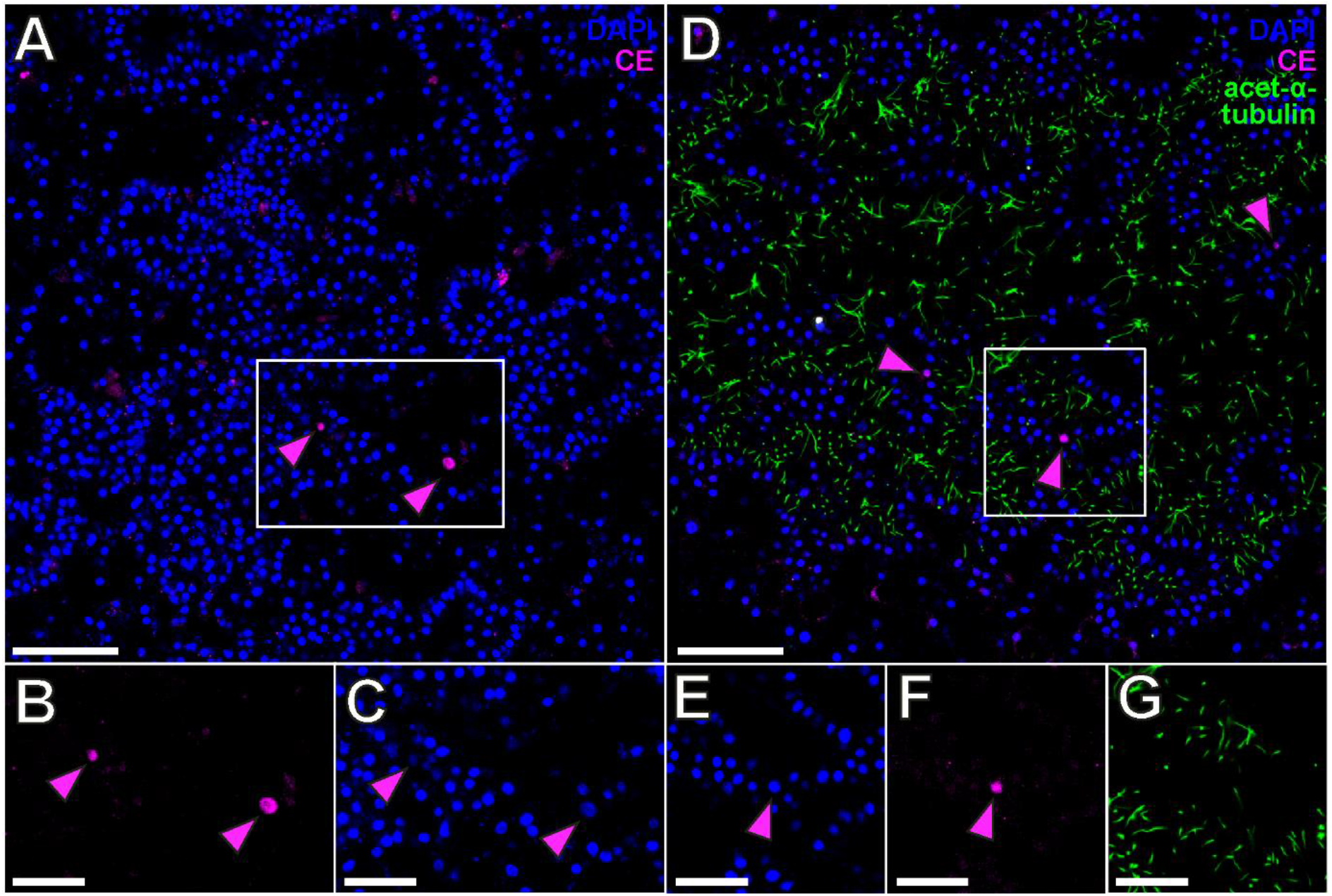
Apoptosis in intact tissues of *Halisarca dujardinii* and *Leucosolenia variabilis*, CLSM. A – the endosome of *H. dujardinii*; B, C – enlarged single-channel images of the zone in the white rectangle in A; D – the proximal part of the oscular tube of *L. variabilis*; E, F, G – enlarged single-channel images of the zone in the white rectangle in D. Several focal planes are combined into one picture. Blue color – DAPI, cell nuclei; magenta – CellEvent, nuclei of apoptotic cells; green – acetylated α-tubulin, flagella. Magenta arrowheads mark CellEvent-positive cells. Scale bars: A, D - 50 μm; B, C, E, F, G - 25 μm.

**Table 4.** Fractions of apoptotic cells in the intact tissues of *H. dujardinii* and *L. variabilis*, CLSM.

| Species | Body region | n | CE-positive cells (% of all cells) |
| --- | --- | --- | --- |
| *Halisarca dujardinii* | Distant from the oscular tube | 7 | 0.03±0.05% |
| *Leucosolenia variabilis* | Oscular tube | 9 | 0.22±0.19% |
|  | Cormus tube | 6 | 0.44±0.66% |
|  | Diverticula | 3 | 0.21±0.16% |

6 Each proportion is given as mean value and SD. Three individuals per species have been examined; n 7 is the number of analyzed stacks.

We also studied suspensions obtained via mechanical dissociation in CMFSW-E. They were stained with CellEvent, fixed, and analyzed via a flow cytometer. Cells of *H. dujardinii* formed a single peak with low/negative CellEvent fluorescence (Figure 10A, B). The fraction of apoptotic cells was low, with CellEvent-positive cells accounting only for 1±1.15% of the total population. In contrast, cells of *L. variabilis* were clearly divided into two subpopulations by the level of CellEvent fluorescence (Figure 10C, D). CellEvent-positive cells with elevated autofluorescence (as evidenced by increased short-wave fluorescence in the PacificBlue channel) comprised 60.8±13.46% of the total cell population (Figure 10C, D). We consider this result as a sampling artifact (see “Discussion” section)

**Figure 10.**
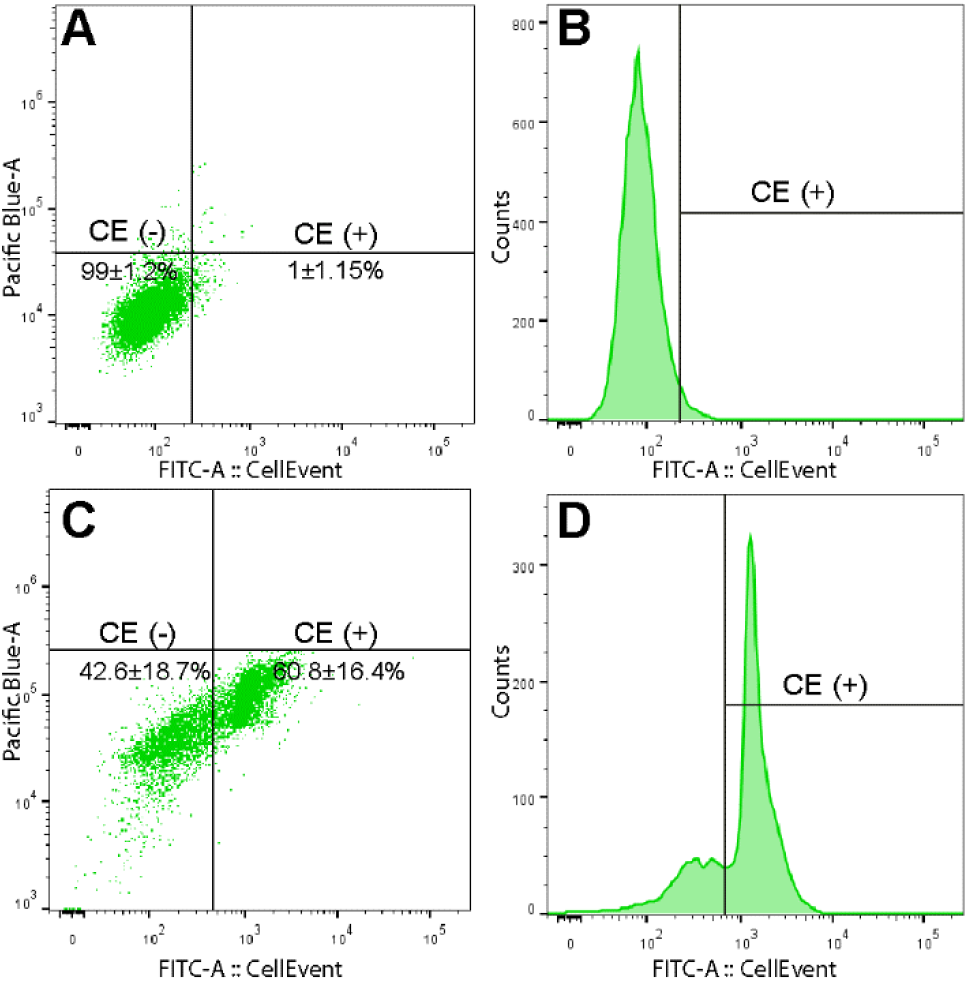
Apoptosis in cell suspensions of *Halisarca dujardinii* (A, B) and *Leucosolenia variabilis* (C, D), flow cytometry. A, C - Pacific Blue-A (autofluorescence) vs FITC-A (CE fluorescence) dot plots; B, D – single-parameter histograms of CellEvent fluorescence. CE(+) population gate is set according to the negative control.

Suspensions were also studied *in vivo* during the experiment, where non-fixed suspensions were stained for 30 min with CellEvent and TMRE (TMRE indicates the presence of the mitochondrial outer membrane potential - i.e., the viability of the mitochondria). These suspensions were studied both by CLSM and flow cytometry. Four cell types were distinguished via CLSM (Figure 11A):

1. TMRE+ CE− cells possessing functional mitochondria (as indicated by TMRE fluorescence) and no active effector caspases (as clear nuclear CellEvent signal is absent);
2. TMRE− CE− cells characterized by the absence of viable mitochondria or active effector caspases (i.e., no CellEvent or TMRE fluorescence observed);
3. TMRE+ CE+ cells possessing both functional mitochondria and active effector caspases (bright TMRE and CellEvent signals are present);
4. TMRE− CE+ cells distinguished by high activity of effector caspases and the loss of the mitochondrial outer membrane potential; these cells display bright nuclear CellEvent signal but no TMRE fluorescence.

**Figure 11.**
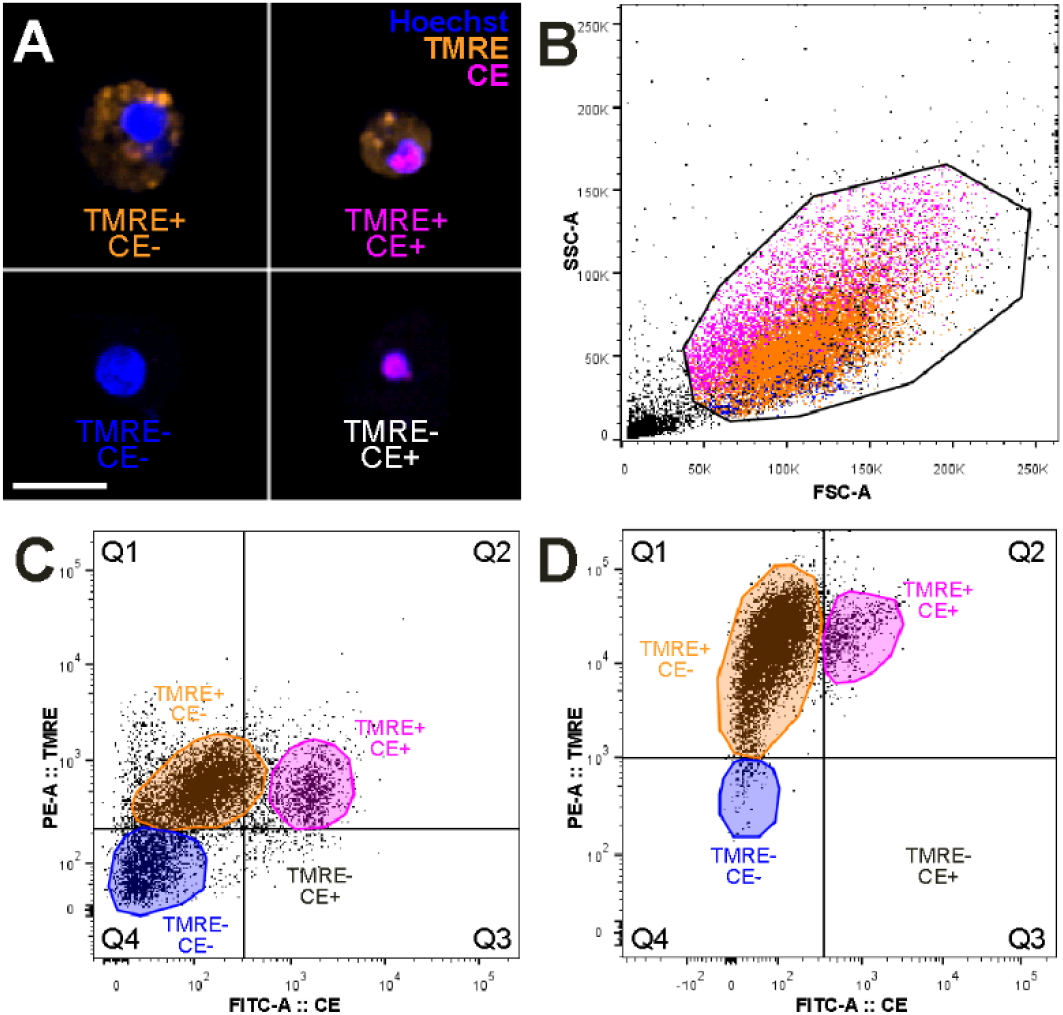
*In vivo* TMRE and CellEvent staining of *Halisarca dujardinii* and *Leucosolenia variabilis* suspensions. A - CLSM of *L. variabilis* suspension, scale bar 10 μm; blue color - Hoechst 33342, cell nuclei; orange - TMRE, functional mitochondria; magenta - CellEvent, nuclei of apoptotic cells; B - scatter plot of FSC-A (cell size) vs SSC-A (cell granularity), an example of primary gating; C, D - TMRE vs CellEvent dot plot of *L. variabilis* (C) and *H. dujardinii* (D) suspensions. The borders of quadrants (Q1-Q4) are set by the negative controls.

The same types of cells were observed using a flow cytometer (Figure 11B-D). Both species demonstrated similar cell distributions. The majority of the cells belonged to the TMRE+ CE− population, which resides in quadrant Q1 and accounted for 89.67±1.06% of the cells in *H. dujardinii* and 70.55±1.48% of the cells in *L. variabilis* (Figure 11C, D; Table 5). The population of the TMRE+ CE+ cells was in the area of higher CellEvent fluorescence, Q2. This population reached 7.51±0.87% in *H. dujardnii* and 8.58±0.77% in *L. variabilis* (Figure 11C, D; Table 5). All TMRE− CE+ cells were in Q3 and do not exceed 1% of the general population and do not form a distinguishable cluster (Figure 11C, D; Table 5). All CellEvent-positive cells (Q2 and Q3) were small and highly granulated, as indicated by light scattering parameters (Figure 11B). Population of TMRE− CE− cells was located in Q4 and accounted for 1.67±0.35% in *H. dujardinii* and 13.70±3.25% in *L. variabilis* (Figure 11C, D; Table 5). While these cells were smaller than TMRE+ CE− cells, they tended to separate from TMRE+ CE+ cells on scatter plots (Figure 11B).

**Table 5.** Fractions of cell populations revealed during the *in vivo* apoptosis investigation, flow cytometry.

| Species | N | TMRE- CE- | TMRE+ CE- | TMRE+ CE+ | TMRE- CE+ |
| --- | --- | --- | --- | --- | --- |
| *Halisarca dujardinii* | 3 | 1.67±0.35% | 89.67±1.06% | 7.51±0.87% | 0 |
| *Leucosolenia variabilis* | 2 | 13.7±3.25% | 70.55±1.48% | 8.58±0.77% | 0.06±0.01% |

- 1 The fraction of each population is represented by mean value and SD. N is the number of analyzed - 2 individuals.

## Discussion

In multicellular animals, the long-term maintenance of tissue homeostasis is achieved by cell turnover^1,4^. Cell turnover is composed of three complementary processes: cell proliferation, cell differentiation and programmed cell death^3^. Sponges are basal metazoans well known for their unique tissue dynamics. In this study, we mainly focused on cell proliferation and apoptosis in intact tissues of sponges, providing promising data on cell turnover in sponges.

### Cell proliferation in intact tissues of Halisarca dujardinii and Leucosolenia variabilis

*Halisarca dujardinii* is a leuconoid demosponge that displays a significant number of cycling cells under the steady-state condition. The presence of EdU- and pH3-positive cells in intact tissue is consistent with flow cytometry data. It should be noted that the distribution of *H. dujardinii* cells by DNA content reveals the classic cell cycle distribution curve^37^. However, the estimates of proliferating cell fractions by CLSM and flow cytometry are slightly different. The proportion of EdU- and pH3-positive cells is about 8% of the total cell number, while S plus G2/M cells together reach 28%. Being undetectable by our CLSM staining, cells in G2-phase could account for the observed difference.

Different regions of the endosome displayed active proliferation and did not differ significantly between the proportions of EdU- and pH3-positive cells. Our results indicate that the proliferation activity is evenly distributed throughout the sponge’s endosome. The oscular tube possesses a different pattern of proliferation and contains few labeled cells. We attribute this difference to the absence of choanocyte chambers and choanocytes in the oscular tube. In other demosponges, choanocytes have been shown to incorporate labeled nucleotides^14,33,34,38^, however, they might have retained the BrdU signal as the progeny of other proliferating cells. Here we demonstrate that choanocytes also make up the majority of cells labeled with anti-pH3 antibodies. Thus, choanocytes are the main proliferating fraction of cells in *H. dujardinii*.

*Leucosolenia variabilis* is a calcareous asconoid sponge that has a similar pattern of cell proliferation as *H. dujardinii*. As in other calcareous sponges, the majority of DNA synthesizing cells are choanocytes^12,38^. Most pH3-labeled cells are also choanocytes, although the proportion of choanocytes among all pH3-labelled cells is lower in *L. variabilis* than in *H. dujardinii*. Body regions containing choanocytes displayed a high number of cycling cells and do not differ in the rate of proliferation. The only exception is in the distal parts of the diverticula. Fewer EdU-positive cells were found there yet the proportion of pH3-positive cells remained nearly the same. This body region also differed in the proportion of choanocytes among the pH3-positive cells, with nearly half of the G2/M cells belonging to the mesohyl/pinacoderm layer. This could be evidence of potential growth since new sclerocytes, pinacocytes and porocytes are obviously needed to lengthen the diverticula. In the oscular rim, which is devoid of choanocytes, the intensity of cell proliferation was very low. There are, however, a few labeled cells, indicating either slow cell turnover or growth.

This study provides the first spatial analysis of cell proliferation across different body regions of sponges representing distinct phylogenetic lines and types of aquiferous systems. In general, proliferation activity appears to be evenly distributed throughout the tissues of sponges (excluding regions that do not contain choanocytes). The proportion of cycling cells in *H. dujardinii* is lower than that of other Demosponges, in which this parameter reaches 20-25% on average^34,38^. Among calcareous sponges, the proportion of cycling cells in *L. variabilis* is slightly higher than in *Sycon coactum*^38^. It should also be noted that during our regeneration studies, the proportion of EdU-labeled cells in *L. variabilis* was overestimated when compared to current data, and is likely an artifact of the counting technique^12^. Similar to the previous studies, we detected that DNA synthesis occurs primarily in the choanocytes; moreover, we show that the distribution of mitotic cells in adult sponges have an overall similar pattern. The proportion of proliferating cells in *H. dujardinii* and *L. variabilis* strongly resembles those animals characterized by fast cell turnover, including platyhelminthes, cnidarians and bivalves^39–42^.

### The contribution of choanocytes to homeostasis

The physiological role of proliferation in intact tissues is either growth or renewal^3^. In order to understand the general dynamics of sponge tissue, we need to figure out the fate of proliferating cells. In intact sponges, cell proliferation takes place in the choanoderm. Since the choanoderm is metabolically active and directly exposed to the environment, we should expect higher rates of choanocyte turnover^4^. Similar to feeding structures of other suspension feeders, constant replacement of old and damaged cells is crucial for the long-term maintenance of the choanoderm^42–45^. Despite constantly facing a potentially hostile environment (e.g., toxic and mutagenic factors dissolved in seawater), choanocytes undergo constant proliferation and represent the main source of new cells in the choanoderm. It should be noted here that the proliferation of morphologically differentiated cells is fairly common among metazoans, including vertebrates^3,46–49^. Cnidarians are particularly interesting given that ecto- and endodermal epithelial cells act not only as barrier and contractile cells, but also retain multipotency and the ability to proliferate^50^.

In fact, the boundary position of choanocytes might be advantageous since cell division requires significant amount of resources. As the primary food-entrapping sponge cells, choanocytes are provided with sufficient energy for constant proliferation and are able to respond appropriately to environmental changes. Moreover, since the choanoderm is obviously enriched by cell-cell interaction, and at least partially compartmentalized by extracellular matrix^18^, it could maintain the proliferative potential of choanocytes through some signaling system resembling that of the stem cell niche. In some animals (e.g., *Hydra* and planarians), the concept of ‘niche’ may be interpreted in a rather broad sense and include a whole organ or tissue as a stem cell niche^69,70^. The choanoderm in sponges might be just an example of such a “tissue niche”.

Since the flagellum is resorbed during cell division, the filtration process becomes impaired. To resolve the conflict between proliferation and food uptake, choanocytes either maintain the proliferating fraction at a certain level, or, minimize the period when the flagellum is absent. The short duration of the M phase is possibly the reason why the proportion of pH3-positive cells does not exceed 1% in any of the examined areas in *H. dujardinii* and *L. variabilis*. Indeed, although the lengths of cell cycle phases vary significantly among different animals, the M phase is always the shortest (e.g., 3-5% of the total duration of the cell cycle in mammals)^51^.

It has been previously shown that choanocytes actively proliferate in intact tissues of demosponges and calcareous sponges. Still, it is usually stated that the proliferation of choanocytes is restricted to the self-renewal of the choanoderm and that the mesohyl contains an autonomous population of cycling cells (e.g., archaeocytes in Demospongiae)^8^. According to Funayama^21^, both choanocytes and certain mesohyl cells represent distinct types of stem cells, however, choanocytes display broad stem cell activity only in calcareous and homoscleromorph sponges^8^. In demosponges and hexactinnelids, archaeocytes are thought to be the active stem cells, with choanocytes being the minor component of the stem cell system, transforming into archaeocytes only under specific circumstances^23^.

However, in both *H. dujardinii* and *L. variabilis* (as well as in several other species), EdU-positive choanocytes outnumber EdU-positive cells in the mesohyl^33,34,38^. Mesohyl cells also account for the minority of the pH3-positive cells. Therefore, mesohyl hardly contains large population of cycling cells. Moreover, it should be taken into account that not all observed EdU-positive cells are actually in the S phase after the prolonged incubation in EdU, since DNA synthesis occurs predominantly in the choanoderm, it is tempting to assume that some of the EdU-positive cells in the mesohyl could actually be choanoderm-derived.

Our EdU tracking experiment in *H. dujardinii* provided us with additional clues. After the removal of the EdU from the medium, the proportion of labeled mesohyl cells continuously increased. There are several ways to interpret this result. It could be that mesohyl cells proliferate much faster than choanocytes, driving the increase in the proportion of labeled cells. However, rapidly cycling archaeocytes would have comprised the majority of labeled cells immediately after the treatment with EdU (i.e., at the initial time point). Thus, there is no evidence that archaeocytes possess short cell cycle. The proportion of labeled mesohyl cells can also increase due to the loss of EdU-positive choanocytes through cell shedding. In several species, choanocytes were found to be massively discarded into the environment^33–35^. Nevertheless, there is little physiological sense in getting rid of newly produced (i.e., EdU-positive) cells, as one would expect that old, non-proliferative choanocytes are those that are predominantly shed. Additionally, extensive cell shedding is characteristic of sponges with high microbial abundance in their tissues^34^. This should not be the case in *H. dujardinii*, which is a low microbial abundance sponge^36^. Moreover, we did not observe extensive cell shedding in this species.

Instead, we suppose that the label flux in *H. dujardinii* occurs via cell migration (in line with similar experiments on planarians^52^). Since choanocytes are the main DNA synthesizing population of cells, the increase in the proportion of EdU-positive mesohyl cells could be a consequence of the migration of choanocytes (or their progeny) into the mesohyl. Of course, the population of archaeocytes might be self-renewing to some extent, but the label flux can be interpreted as indirect evidence that choanocytes contribute to the pool of mesohyl cells. Even if archaeocytes take part in the renewal of the choanoderm, the general tissue dynamics leans towards the opposite direction. The same process may occur in intact tissues of *L. variabilis*. Although we have not performed pulse-chase experiments in this species, we hypothesize that the population of amoeboid mesohyl cells in this species is even more transient and choanocyte-dependent.

Therefore, choanocytes are apparently the major part of cycling cells not only in the calcareous sponge *L. variabilis*, but also in the demosponge *H. dujardinii*. It seems that the proliferation of choanocytes is not only limited to the renewal of the choanoderm, as it has been previously shown that choanocytes contribute to gametogenesis^8,53–55^, regeneration^12,14,16^ and can express certain stem cell markers (at least in demosponges and homoscleromphs)^26,28,56,57^. Additionally, it seems that choanocytes may also participate in cell turnover through cell migration. Together, this data suggests that the role of choanocytes in the stem cell system of sponges is underestimated. Choanocytes might act as a major source of new cells (including cells of mesohyl) in both calcareous sponges and demosponges during regular cell turnover or growth.

### Apoptotic activity in intact tissues of sponges

Under the steady-state condition, cell proliferation is counterbalanced by the process of cell elimination. Apoptosis is considered to be the most common way of cell death in animals^3,32,58^. It is a tightly regulated and complex process resulting in the fragmentation of a dying cell into apoptotic bodies^31^. Once these apoptotic bodies are engulfed by other cells, neat cell death is reached. It should be noted that tissues exposed to adverse environments usually display high rates of physiological turnover (e.g., mammalian epidermis and intestine; gills, intestine and mantles of mollusks; epithelia of ascidians and cnidarians)^42,58–62^. Given that choanocytes are highly proliferative, we would expect a significant amount of dying cells.

CLSM studies using CellEvent staining indicated that apoptotic rate is rather low in intact tissues of *H. dujardinii* and *L. variabilis*, with less than 1% of cells containing active effector caspases. We consider this result to be the most relevant, as it fits well with other known studies on sponges^16,33,34^. Insignificant number of apoptotic cells was initially thought to indicate the possibility of caspase-independent cell death in intact sponges. However, low number of apoptotic cells is also characteristic of tissues of other animals under the steady-state condition^63,64^. Thus, we suggest that apoptosis is involved in regular tissue renewal in sponges.

Significantly more CellEvent-positive cells were found in the CMFSW-E suspensions analyzed using flow cytometry. These cells are small, highly granulated and contain active effector caspases, thus they demonstrate the classic features of apoptosis^65,66^. Suspensions of *H. dujardinii* contained few apoptotic cells, whereas CellEvent-positive cells in *L. variabilis* comprised up to 60% of the general population. SubG1 cells detected by means of DNA content are also important to mention here. These small, highly granulated cells characterized by fragmented DNA should be considered apoptotic. Since cell debris and bacteria might also be detected as a SubG1 population, it does not represent an accurate estimation of the number of apoptotic cells. However, SubG1 cells may serve as indirect and rough evaluation of the apoptotic activity. The presence of SubG1 cells seems to be an inevitable consequence of our sample preparation^67^. Yet, the proportion of SubG1 cells differ drastically between species; CMFSW-E dissociation does not significantly affect *H. dujardinii*, but leads to a massive transition of *L. variabilis* cells to the SubG1 population. Obviously, CMFSW-E suspensions of *L. variabilis* indicate massive cell death caused by sample preparation, where the apoptotic response develops quickly within one or two hours. More importantly, this was not the case in suspensions made using pre-fixed tissues. It seems that dissociation results in the activation of the apoptotic machinery in cells of *L. variabilis*. Thus, the observed number of CellEvent-positive cells and SubG1-cells in the suspensions of *L. variabilis* cannot be considered an adequate estimation of the overall apoptotic activity.

The observed interspecific variability of the apoptotic response might be associated with the phylogenetic position and type of body organization. The majority of the body in *H. dujardinii* (Demospongiae) is occupied by mesenchymal tissue, and in comparison, *L. variabilis* (Calcarea) is mainly composed of epithelial-like layers of choanoderm and pinacoderm. Overall, it appears that calcareous sponges have a more epithelial-like organization, relying on tighter cell-to-cell interactions. For example, regeneration occurs via blastema formation and mesenchymal morphogeneses in demosponges, while in calcareous sponges, it depends on the morphallactic reorganization of adjacent intact tissue through epithelial morphogeneses^11,12,68^. The disruption of the natural system of cell interactions during the CMFSW-E dissociation could lead to the massive induction of apoptotic activity in calcareous sponges.

Certainly, some degree of apoptosis upregulation should be expected in non-fixed suspensions. Apoptotic cells accounted for 7-8% of the general population in both species during *in vivo* studies; however, despite the apoptosis upregulation, *in vivo* experiments shed some light on the mechanisms of apoptosis. In mammalian cells, double staining with TMRE and CellEvent usually reveals the presence of three populations: 1) intact TMRE+ CE− cells characterized by intact mitochondria; 2) early apoptotic TMRE− CE− cells, in which mitochondrial transmembrane potential has vanished; and 3) late apoptotic TMRE− CE+ cells that contain non-functioning mitochondria and active effector caspases^69^. This situation is typical for mammalian cells since the permeabilization of the inner mitochondrial membrane is shown to be tightly associated with the mitochondrial outer membrane permeabilization (MOMP)^63^. In turn, MOMP is the critical event during the apoptotic response, resulting in the release of some pro-apoptotic proteins from the mitochondrial intermembrane space into the cytoplasm. The most crucial pro-apoptotic protein is cytochrome c, which is able to bind to cytoplasmic protein APAF1, forming the apoptosome and beginning to recruit effector caspases^32,70^. In flatworms (*Dugesia dorotocephala* and *Schistosoma mansonii*) and echinoderms (*Strongylocentrotus purpuratus* and *Dendraster excentricus*), the activation of effector caspases is also cytochrome-c dependent^71^. Thus, active effector caspases in their cells should be detected only after the inner membrane permeabilization and MOMP. Some invertebrates, however, do not fit into this scenario^72^. In common model organisms like *Caenorhabditis elegans* and *Drosophila melanogaster*, the activation of effector caspases is not associated with MOMP. In such cases, the promotion of apoptotic cell death does not depend on the presence of cytochrome c in the cytoplasm, even if MOMP is present, it is more of a consequence than a reason for apoptotic response^73,74^.

Our data indicates that apoptosis in sponges might resemble the scenario of *C. elegans* and *D. melanogaster*, as significant amounts of TMRE+ CE+ cells have been observed in *H. dujardinii* and *L. variabilis* suspensions. TMRE+ CE+ cells contain active effector caspases but possess well-functioning mitochondria. Unstained negative control suspensions indicated that the high level of TMRE fluorescence in these presumably early apoptotic cells is specific: cells of both *H. dujardinii* and *L. variabilis* maintained functional mitochondria during apoptosis. Thus, there is a possibility that the apoptosis in sponge cells is MOMP-independent. Therefore, and in contrast to mammalian cells, the TMRE− CE− population seen in these two sponges may contain not early apoptotic cells as originally thought, but represents a population of damaged or dead cells and cell debris. Rare TMRE− CE+ cells might be interpreted as non-viable, late apoptotic cells with non-functional mitochondria.

### Conclusions

Here we provide quantitative and spatial analysis of cell proliferation and apoptosis in two sponges belonging to different classes of Porifera. Intact tissues of *Halisarca dujardinii* (Demospongiae) and *Leucosolenia variabilis* (Calcarea) are highly proliferative, indicating either high rate of growth or cell turnover. In both species, the most proliferating fraction of cells are choanocytes, with proliferation occurring evenly throughout areas of the body containing these cells. The proliferation of choanocytes maintains tissue homeostasis in the choanoderm, and possibly, in the mesohyl/pinacoderm. Further investigations with *in vivo* cell tracking and *in situ* hybridization are needed to prove if the choanocytes, or their progeny, actively differentiate into other cell types. The number of apoptotic cells in intact tissues of both species is low, although it corresponds with the data on other animals. Therefore, we still think the apoptosis is involved in the maintenance of tissue homeostasis.

Our data will serve as a basis for further detailed research of tissue homeostasis in sponges. While we have an information about spatial distribution and intensity of cell proliferation and apoptosis for a number of sponge species, many other processes and mechanisms contributing to tissue homeostasis in sponges remains uncharacterized. Behavior and fate of cell progeny, temporal characteristics of the proliferation, apoptosis and differentiation, and, finally, the signaling systems maintaining proliferative potential of choanocytes are the aims of the future study. Altogether, this data would help us to establish a comprehensive model of tissue dynamics in these basal metazoans.

## Experimental Procedures

### Sampling and laboratory maintenance

*Halisarca dujardinii* (Demospongiae, Chondrillida) and *Leucosolenia variabilis* (Calcarea, Leucosolenida) were collected in the Kandalaksha Bay in the White Sea near the Pertsov White Sea Biological Station of Lomonosov Moscow State University (66°34′ N, 33°08′ E) in August 2019 and 2020. Sponges were collected in the upper subtidal zone with the substrate (brown algae *Fucus* and *Ascophyllum*) and cleared of epifauna. Specimens were kept in laboratory aquariums with running natural seawater of ambient temperature. Each experiment was carried out on individuals of the same size collected on the same day. Sponges were kept in open plastic Petri dishes filled with 5 mL of 0.22 μm-filtered seawater (FSW) during the experiments. Petri dishes were kept between 8-12 °C on a shaker for better stirring and aeration of FSW.

### Histological studies of intact tissues

For histological studies, sponge tissues were fixed for 2 hours at 4 °C with 2.5% glutaraldehyde (Electron Microscopy Science, 16020) and post-fixed for 1 hour at room temperature (RT) with 1% OsO_4_ (Electron Microscopy Science, 19100). Fixation and post-fixation were done in modified 0.1M Na-Cacodylate buffer (0.1M Na-Cacodylate, 85.55 mM NaCl, 5 mM CaCl_2_, 5 mM MgCl_2_; pH 7.0-7.5). After the post-fixation, specimens were dehydrated and embedded in Epon/Araldite epoxy embedding media (Electron Microscopy Science, 13940) according to the standard protocol previously described^12,75^.

Semi-thin sections (1 μm) were cut using LKB V and Leica UC6 ultramicrotomes. Sections were stained with 1% toluidine blue – 0.2% methylene blue mixture for 1-1.5 min at 60 °C and examined with a Carl Zeiss Axioplan 2 microscope (Carl Zeiss) equipped with an AxioCam HRm (Carl Zeiss) digital camera and AxioVision 3.1 (Carl Zeiss) software.

### Study of cell proliferation with CLSM

We used 5-ethynyl-2’-deoxyuridine (EdU) and Ser10-phosphorylated histone 3 (pH3) monoclonal antibodies as the markers of cell proliferation. EdU is a nucleoside analog of thymidine, which incorporates into nuclei of proliferating cells during DNA synthesis, marking cells in the S-phase^76^. Antibodies were used against phosphorylated histone 3 labeled cells in the late G2-phase and M-phase^77^. To mark the flagella of the choanocytes, we used antibodies against acetylated-α-tubulin.

The EdU (Lumiprobe, 10540) stock solution was prepared in dimethyl sulfoxide (DMSO, MP Biomedicals 196055). Optimal EdU concentration and incubation times were discovered during preliminary studies. *Halisarca dujardinii* individuals were incubated in 5 mL of FSW with 200 μM EdU for 12 hours and individuals of *L. variabilis* were incubated in 5 mL of FSW with 20 μM EdU for 6 hours. Sponges kept in 5 mL FSW with 100 or 10 μM DMSO served as negative controls, respectively. We examined eight individuals of *H. dujardinii* and six individuals of *L. variabilis*.

After the EdU incubation, all individuals were fixed with 4% paraformaldehyde in phosphate-buffered saline (4% PFA PBS, Carl Roth 0335.2) for 12 hours at 4 °C. Fixed specimens were rinsed with PBS several times, then they were incubated in 1-3% bovine serum albumin in PBS (BSA, MP Biomedicals 0216006980) and permeabilized using 0.5% Triton X-100 (Sigma-Aldrich T8787) in PBS. Click-reaction (visualization of EdU) was performed for 1 hour at RT in a solution of 4 mM CuSO_4_, 10 μM Sulfo-Cyanine3 Azide (Lumiprobe A1330) and 20 mg/mL sodium L-ascorbate (Sigma-Aldrich 11140) in PBS. Click-reaction and subsequent manipulations were carried out in the dark. Specimens were blocked with a blocking solution of 1% BSA, 0.1% gelatin from cold-water fish skin (Sigma-Aldrich G7041), 0.5% Triton X-100 and 0.05% Tween-20 (Sigma-Aldrich P1379) in PBS and then incubated in the primary antibodies (Rabbit Anti-phospho-histone H3, Sigma-Aldrich H0412 and Mouse Anti-acetylated-α-tubulin, Sigma-Aldrich T6793, both diluted 1:1000 in blocking solution) overnight at 4 °C. Afterwards, specimens were rinsed twice with the blocking solution and treated with the secondary antibodies (Donkey anti-Mouse IgG Alexa Fluor 488, ThermoFisher Scientific A21202 and Donkey anti-Rabbit IgG Alexa Fluor 647, ThermoFisher Scientific A-31573 in blocking solution, both with a final concentration 1 μg/mL) overnight at 4 °C. Cell nuclei were stained with 4′,6-diamidino-2-phenylindole (DAPI, Acros 202710100) in PBS (final concentration 2 μg/mL). Spicules of *L. variabilis* were dissolved using 5% ethylenediaminetetraacetic acid (EDTA) in distilled water for 30 min at RT. Finally, specimens rinsed with PBS were mounted in 90% glycerol (MP Biomedicals 193996) with 2.5% 1,4-diazabicyclo[2.2.2]octane (DABCO, Sigma-Aldrich D27802) and studied using a confocal laser scanning microscope.

Cell proliferation was studied in several body regions of the sponges. Three regions were studied in *H. dujardinii*: 1) the endosome adjacent to an oscular tube (n=8); 2) the endosome at some distance (approximately half the body radius) from an oscular tube (n=8); and 3) the oscular tube (n=4). Seven regions were studied in *L. variabilis*: 1) the oscular rim (n=10); 2) the distal part of the oscular tube (n=10); 3) the middle part of the oscular tube (n=10); 4) the proximal part of the oscular tube (n=10); 5) the distal part of the diverticulum (n=14); 6) the proximal part of the diverticulum (n=14); and 7) the cormus tube (n=11). Studied fields of view within each region were chosen randomly. For subsequent analysis, Z-stacks were obtained from each field of view.

Z-stacks were obtained by CLSM (Nikon A1, Nikon, Shinagawa, Japan) using lasers of wavelength 405 nm (DAPI), 488 nm (Mouse Anti-acetylated-α-tubulin ABI + DAM IgG Alexa Fluor 488 ABII), 561 nm (EdU + Sulfo-Cyanine3 Azide) and 648 nm (Rabbit Anti-phospho-histone H3 ABI + DAR IgG Alexa Fluor 647). All Z-stacks were obtained with a 1-μm Z-step, 48 μm (*H. dujardinii*) and 68 (*L. variabilis*) μm thick on average.

During subsequent Z-stack analysis, we estimated the total fraction of EdU- and pH3-positive nuclei (as a proportion of EdU-/pH3-positive nuclei to all nuclei stained with DAPI), the fractions of EdU- and pH3-positive choanocytes and mesohyl cells (as a proportion of EdU-/pH3-positive choanocytes/mesohyl cells to all nuclei stained with DAPI) and the fraction of choanocytes among all EdU/pH3-positive cells (as a proportion of EdU-/pH3-positive choanocytes to all EdU-/pH3-positive cells).

Z-stack analysis was performed using Bitplane Imaris v7.2.1 software. Each stack of *H. dujardinii* and *L. variabilis* contained 6000-7000 and 2000-4000 cells, respectively. Choanocytes were identified by the presence of flagella and the characteristic shape of the choanoderm/choanocyte chambers. Mesohyl cells were separated from choanocytes via the «Surface» tool. Cells of interest (e.g., EdU- or pH3-positive cells) were counted by the «Spots» tool. We combined automatic identification of nuclei via Quality/Intensity Median thresholds with manual correction of spots. This counting procedure implies 3D visualization of an entire Z-stack and allows us to precisely distinguish between labeled cells of choanoderm and mesohyl.

### EdU tracking in Halisarca dujardinii

Intact *H. dujardinii* individuals were incubated in 5 mL of FSW with 200 μM EdU for 12 hours and then transferred into a running seawater aquarium. Sponges incubated in 5 mL of FSW with 100 μL of DMSO were used as negative controls. Sponges were fixed with 4% PFA PBS on day 1, 5 and 7 after being transferred into aquariums (3, 3 and 5 individuals, respectively). Individuals fixed immediately after the EdU incubation served as the initial time point, day 0. All specimens were treated for CLSM studies as described above (see “Study of cell proliferation with CLSM” section), but no antibody staining was performed in this experiment. We studied areas of the endosome at some distance from an oscular tube. One randomly chosen field of view within the region was analyzed in each individual. For subsequent analyses, a Z-stack was obtained from each field of view. We estimated the fractions of choanocytes and mesohyl cells among all EdU-positive cells (as a proportion of EdU-positive choanocytes/mesohyl cells to all EdU-positive cells). As a result, the main parameter of the analysis was the mean proportion of choanocytes and mesohyl cells in the total number of EdU-positive cells. Details of data acquisition, nuclei counting and further analysis have been described above (see “Study of cell proliferation with CLSM” section). In this experiment, choanocyte chambers were identified by their characteristic shape visualized by DAPI staining.

### Evaluation of cell cycle distribution

We combined calcium and magnesium-free seawater (20 mM KCl, 300 mM NaCl, 10 mM Tris-HCl and 15 mM EDTA in distilled water) (CMFSW-E) and mechanical tissue dissociation to obtain single-cell suspensions. Sponges were cleaned of epifauna, soaked in CMFSW-E for 30 min and squeezed into CMFSW-E through a sterile gauze with a mesh size of 30 μm. After 30 min, suspensions were centrifuged and resuspended in a new aliquot of CMFSW-E. After another 30 min, suspensions were fixed with 70% cold ethanol at −20 °C. Unfortunately, this dissociation technique yielded a poor-quality cell cycle curve in *L. variabilis* and an alternative fixation method was required. The new dissociation technique used intact pieces of *L. variabilis*, which were first fixed in cold 70% ethanol. After the fixation, tissues were soaked in CMFSW-E and mechanically dissociated through gauze with a mesh size of 30 μm.

Cell suspensions from five specimens of *H. dujardinii* and five specimens of *L. variabilis* were studied for cell cycle distribution, using propidium-iodide (PI) staining since PI fluorescence is proportional to DNA content^37^. Cell suspensions were resuspended in PBS and stained with 1 μg/mL PI (P1304MP, ThermoFisher Scientific) for at least 12 hours. Unstained suspensions served as a negative control. Suspensions were analyzed by a FACSAria SORP instrument (BD Biosciences, USA) using an excitation laser wavelength of 561 nm (PI excitation); 50 000 events were collected per sample. All collected data was processed using FlowJo V10. Primary gating was performed according to forward (FCS-A) versus side scatter (SSC-A) parameters; secondary gating on singlet cells according to FSC-W versus FSC-H parameters.

To obtain cell cycle distributions, we used the internal FlowJo CellCycle plugin. The Watson model approximates the DNA content curve as the sum of two Gaussian distributions represented by G0/G1 and G2/M cells, considering cells between them as cells in the S phase. In most cases, standard parameters were used. When the G2/M peak was hard to distinguish, we constrained its position depending on the relative positions of the G0/G1 and G2/M peaks. No CV constraint was performed.

### Investigation of apoptotic activity in intact tissues

Tetramethylrhodamine ethyl ester (TMRE, ThermoFisher Scientific, T669) and CellEvent Caspase-3/7 Green ReadyProbes Reagent (ThermoFisher Scientific, R37111) (CellEvent) were used as markers of apoptotic activity for both CLSM and flow cytometry studies. The absence of TMRE fluorescence serves as a sign of mitochondrial inner membrane permeabilization, which is characteristic of the early stages of apoptosis. CellEvent is an indicator of activated effector caspases present in cells during the middle and late stages of apoptosis^63^. In each experiment, we used unstained specimens as negative controls.

#### CellEvent staining of intact tissues for CLSM

Specimens (3 sponges per species) were incubated in 1 mL of FSW with CellEvent (dilution 1:100) for 2 hours at 10 °C in the dark. After the incubation they were fixed with 4% PFA PBS. Then specimens were fixed with 4% PFA PBS, washed with PBS and blocking solution for several hours, and stained with DAPI as described above (see “Study of cell proliferation with CLSM” section). Specimens of *L. variabilis* were stained with acetylated-α-tubulin antibodies to identify choanocytes. In *H. dujardinii*, no antibody staining was performed and choanocyte chambers were identified by characteristic shape of choanocyte chambers visualized by DAPI staining. During this experiment, we used lasers of 405 nm (DAPI), 488 nm (CellEvent) and 648 nm (Mouse Anti-acetylated-α-tubulin ABI + DAM IgG Alexa Fluor 647 ABII) wavelengths. Details of data acquisition and nuclei counting are provided in the “Study of cell proliferation with CLSM” section. Any nuclei displaying bright CellEvent staining was considered apoptotic. During the analysis, we estimated the total fraction of CellEvent-positive cells (as a proportion of CellEvent-positive nuclei to all nuclei labeled with DAPI) and the proportion of choanocytes/mesohyl cells among all CellEvent-positive cells (as a proportion of CellEvent-positive choanocytes/mesohyl cells to all CellEvent-positive cells).

#### CellEvent staining of suspensions for flow cytometry

We took samples (250 mL) containing 1.5×10^6^ cells from non-fixed CMFSW-E suspensions of *H. dujardinii* and *L. variabilis* (4 samples from 4 sponges per species). The suspensions were obtained as described in the “Evaluation of cell cycle distribution” section. The samples were incubated in CellEvent (dilution 1:100) at 10 °C for 30 min in the dark. Samples were then centrifuged and fixed with 4% PFA PBS at 4 °C and stored for 2 months. Prior to analyses by flow cytometry, cells were washed three times with PBS. All suspensions were analyzed by a FACSAria SORP instrument (BD Biosciences, USA) using an excitation laser wavelength of 488 nm (CellEvent excitation). Primary gating was as described above (see the “Evaluation of cell cycle distribution in intact tissues” section); the final quadrant gating to distinguish CellEvent-positive cells was set according to the unstained control. Any event exceeding the autofluorescence of the unstained control was considered a CellEvent-positive (apoptotic) cell. Subsequent flow cytometry studies showed that *L. variabilis* suspensions contained an anomaly high number of CellEvent-positive (apoptotic) cells. Thus, the data obtained from these samples were not used in the main analysis.

#### Vital staining of suspensions for CLSM and flow cytometry

For vital CLSM studies, the suspensions were obtained as described in the “Evaluation of cell cycle distribution” section. Suspensions were stained with TMRE (final concentration - 100 μM), CellEvent (dilution 1:100) and Hoechst 33342 (dilution 1:100) at 10 °C for 30 min in the dark. During this experiment, we used lasers of wavelength 405 nm (DAPI), 488 nm (CellEvent) and 561 nm (TMRE). Data acquisition was the same as previously described above (see “Study of cell proliferation with CLSM” section). No quantitative analysis was performed; acquired images were used to illustrate flow cytometry data.

For the vital flow cytometry study, three individuals of *H. dujardinii* (August 2020) and two individuals of *L. variabilis* (February 2020) were transported to Moscow in thermoses filled with natural seawater at 10 °C. Sponges were dissociated in CMFSW-E for 30 min and stained with TMRE and CellEvent as previously described. After staining, the suspensions were analyzed using a FACSAria SORP instrument (BD Biosciences, USA) with an excitation laser wavelength of 488 nm (CellEvent excitation) and 561 nm (TMRE excitation). Primary gating was as described above (see the “Evaluation of cell cycle distribution in intact tissues” section). The final quadrant gating to distinguish between different cell populations was performed according to the unstained control. Any event exceeding the autofluorescence of the unstained control was considered a TMRE/CellEvent-positive cell.

### Statistical analysis and plotting

Statistical analysis and plotting were performed in GraphPad PRISM v8.0.1. Mean values and standard deviations were calculated for every parameter (the proportions of EdU-positive cells, pH3-positive cells, etc.). To compare these parameters between different body parts or time points, we used Kruskal-Walles test followed by Dunn test since it is a distribution-free test appropriate for multiple comparisons between small data subsets. The significance level was 0.05. All p-values represented in the text are Dunn test p-values. Tables with exact p-values for all tests are provided in the Mendeley Data root directory (see “Availability of data and material” section).

## Acknowledgments

The authors acknowledge the support of Lomonosov Moscow State University Program of Development (FACSAria SORP flow cytometer/sorter, Nikon A1 CLSM) and Center of microscopy WSBS MSU. Authors sincerely thank Darya Potashnikova (Lomonosov Moscow State University) for operating the flow cytometer, Daniyal Saidov (Lomonosov Moscow State University), for statistical analysis tips, Vitaly Kozin and Alexandra Shalaeva (Saint-Petersburg State University) for helping with Bitplane Imaris, Olga Vorobjeva (Lomonosov Moscow State University), for collection and transporting *L. variabilis* individuals for the *in vivo* apoptosis investigation, Igor Kosevich (Lomonosov Moscow State University) and Ilya Borisenko (Saint-Petersburg State University) for helpful tips and advice. Authors also thanks Brett C. Gonzales (Smithsonian National Museum of Natural History) for his invaluable help with English language corrections.

## Authors’ contributions

NM, AL and AS designed the study. NM, AL, FB, KS and VF collected the material, carried out CLSM proliferation studies and analyzed data in Bitplane Imaris. NM conducted experiments and analyzed data from the pulse-chase EdU studies, apoptosis investigations in intact tissues, DNA content and apoptosis evaluation in cell suspensions, and performed the statistical analysis. AL, NM, AE and AS prepared the manuscript with contributions from all authors. All authors reviewed and approved the final manuscript.

## Availability of data and materials

The draft dataset supporting the conclusions of this article is available in the Mendeley Data repository, https://data.mendeley.com/datasets/hhmc8t6gv7/draft?a=ee1a45f9-2b2d-4b65-b863-ec4e84785263 (*H.dujardinii;* doi:10.17632/hhmc8t6gv7.1) and https://data.mendeley.com/datasets/vbk6h8vcyy/draft?a=97e2ee7f-68e0-455f-b36b-2ad25872ff9a (*L. variabilis*; doi:10.17632/vbk6h8vcyy.1).

## List of abbreviations

FSW: filtered seawater
EdU: 5-ethynyl-2’-deoxyuridine
pH3: Ser10-phosphorylated histone 3
DMSO: dimethyl sulfoxide
PFA: paraformaldehyde
PBS: phosphate-buffered saline
CMFSW-E: calcium and magnesium-free seawater
TMRE: tetramethylrhodamine ethyl ester
CellEvent or CЕ: Caspase-3/7 Green ReadyProbes™ Reagent
PI: propidium iodide
CLSM: confocal laser scanning microscopy
MOMP: mitochondrial outer membrane permeabilization

